# Immune Responses in Checkpoint Myocarditis Across Heart, Blood, and Tumor

**DOI:** 10.1101/2023.09.15.557794

**Authors:** Steven M. Blum, Daniel A. Zlotoff, Neal P. Smith, Isabela J. Kernin, Swetha Ramesh, Leyre Zubiri, Joshua Caplin, Nandini Samanta, Sidney C. Martin, Alice Tirard, Pritha Sen, Yuhui Song, Jaimie Barth, Kamil Slowikowski, Mazen Nasrallah, Jessica Tantivit, Kasidet Manakongtreecheep, Benjamin Y. Arnold, John McGuire, Christopher J. Pinto, Daniel McLoughlin, Monica Jackson, PuiYee Chan, Aleigha Lawless, Tatyana Sharova, Linda T. Nieman, Justin F. Gainor, Dejan Juric, Mari Mino-Kenudsen, Ryan J. Sullivan, Genevieve M. Boland, James R. Stone, Molly F. Thomas, Tomas G. Neilan, Kerry L. Reynolds, Alexandra-Chloé Villani

**Affiliations:** Center for Immunology and Inflammatory Diseases, Department of Medicine, Massachusetts General Hospital, Boston, MA, USA; Massachusetts General Hospital, Cancer Center, Boston, MA, USA; Broad Institute of Massachusetts Institute of Technology and Harvard, Cambridge, MA, USA; Harvard Medical School, Boston, MA, USA; Cardio-Oncology Program, Division of Cardiology, Department of Medicine, Massachusetts General Hospital, Boston, Massachusetts, USA; Transplant and Immunocompromised Host Program, Division of Infectious Diseases, Department of Medicine, Brigham and Women’s Hospital; Department of Pathology, Massachusetts General Hospital, Boston, MA, USA; Division of Rheumatology, North Shore Physicians Group, Department of Medicine, Mass General Brigham Healthcare Center, Lynn, MA, USA; Clinical Research Center, Massachusetts General Hospital, Boston, MA, USA; Department of Surgery, Massachusetts General Hospital, Boston, MA, USA; Division of Gastroenterology, Department of Medicine, Massachusetts General Hospital, Boston, Massachusetts, USA

**Author notes:** These authors contributed equally (Steven M. Blum, Daniel A. Zlotoff, Neal P. Smith, Isabela J. Kernin, Swetha Ramesh). Co-senior authors (Molly F. Thomas, Tomas G. Neilan, Kerry L. Reynolds, Alexandra-Chloé Villani). Corresponding author: Alexandra-Chloé Villani, PhD Massachusetts General Hospital, 149 13th Street, Room 8102 Charlestown, MA 02129.

## Abstract

Immune checkpoint inhibitors (ICIs) are widely used anti-cancer therapies that can cause morbid and potentially fatal immune-related adverse events (irAEs). ICI-related myocarditis (irMyocarditis) is uncommon but has the highest mortality of any irAE. The pathogenesis of irMyocarditis and its relationship to anti-tumor immunity remain poorly understood. We sought to define immune responses in heart, tumor, and blood during irMyocarditis and identify biomarkers of clinical severity by leveraging single-cell (sc)RNA-seq coupled with T cell receptor (TCR) sequencing, microscopy, and proteomics analysis of 28 irMyocarditis patients and 23 controls. Our analysis of 284,360 cells from heart and blood specimens identified cytotoxic T cells, inflammatory macrophages, conventional dendritic cells (cDCs), and fibroblasts enriched in irMyocarditis heart tissue. Additionally, potentially targetable, pro-inflammatory transcriptional programs were upregulated across multiple cell types. TCR clones enriched in heart and paired tumor tissue were largely non-overlapping, suggesting distinct T cell responses within these tissues. We also identify the presence of cardiac-expanded TCRs in a circulating, cycling CD8 T cell population as a novel peripheral biomarker of fatality. Collectively, these findings highlight critical biology driving irMyocarditis and putative biomarkers for therapeutic intervention.

## Introduction

Immune checkpoint inhibitors (ICIs) improve cancer outcomes for a wide range of tumor types^1^ by blocking critical negative regulatory signals in the immune system, but their disruption of peripheral tolerance can cause immune-related adverse events (irAEs) that can affect nearly every organ system^2^. Severe irAEs may lead to ICI treatment termination, morbidity, lasting disability, and death^3,4^. ICI-related Myocarditis (irMyocarditis) occurs in 0.3-1.7% of ICI recipients^5–8^ yet carries a mortality rate of 20-50%, which is the highest of any irAE and 10-fold higher than myocarditis from other causes^9–11^. Like most irAEs, the molecular underpinnings of irMyocarditis remain poorly understood, which has hampered efforts to design effective risk-stratification, diagnostic, and therapeutic strategies. Diagnosis is challenging and commonly requires an endomyocardial biopsy^12^, which is not widely available in most clinical settings and carries significant risks^13^. As there is a lack of prospective therapeutic clinical trial data to inform the management of irMyocarditis, treatment is often guided by expert consensus and extrapolation of data for other conditions, such as cardiac allograft rejection^14^. Emerging non-randomized data has supported the use of more tailored immunosuppressive strategies^15^. However, some immunosuppressive agents, such and corticosteroids and JAK-STAT inhibitors, may be associated with other adverse cardiovascular outcomes and have the potential to counteract anti-tumor immune responses^16–18^.

irMyocarditis is histologically characterized by a patchy, heterogeneous lymphohistiocytic infiltrate in the myocardium, which includes both CD4^+^ and CD8^+^ T cell infiltrates^7,8,19,20^. Furthermore, shared T-cell clones were reported in paired myocardium, skeletal muscle, and tumor in a patient with irMyocarditis and ICI-related myositis (irMyositis)^19^. Bulk RNA sequencing analysis of myocardial tissue from patients with irMyocarditis demonstrated upregulation of many interferon-stimulated genes (ISGs), including *CXCL9*, *MDK*, and *GBP5*^21^. Furthermore, irMyocarditis patients show clonal expansion of circulating CD8^+^ T cells expressing cytotoxic genes and high levels of inflammatory chemokines^22^. Studies in *Pdcd1^−/−^Ctla4^+/–^* mice, a genetic model of irMyocarditis, demonstrated that effector and proliferating CD8^+^ T cells in the myocardium were necessary and sufficient to elicit myocarditis^23,24^, though the fidelity of mouse model systems to human irMyocarditis remains unclear^24–28^.

In this study, we aimed to investigate four poorly understood aspects of irMyocarditis pathophysiology, which may overlap with other irAEs: (1) the cellular microenvironment in irMyocarditis heart tissue that may drive and sustain disease pathogenesis; (2) the clonal and phenotypic relationship between immune cells in the heart and blood; (3) the degree of T-cell clonal sharing between the heart and paired tumors; and (4) the identity of biomarker candidates that could stratify patients with potentially more severe or fatal irMyocarditis. We leveraged single-cell RNA-sequencing (scRNA-seq) and paired T cell receptor (TCR) sequencing of heart tissue, blood, and tumor to define the cellular phenotype and molecular transcriptional programs associated with irMyocarditis. We found that the myocardial microenvironment is enriched with multiple immune subsets and fibroblasts that upregulate inflammatory genes whose protein products could potentially be targeted by existing immunosuppressive medications. Our study is the first to demonstrate that T cell clonotypes enriched in tumors and heart tissue from irMyocarditis patients are largely distinct. Finally, we identify candidate biomarkers of disease onset, cardiac injury, and fatal irMyocarditis. These findings reveal the cellular and transcriptional changes that underlie irMyocarditis and enable the identification of putative targets for clinical diagnostics and tailored clinical intervention.

## Results

### Study Design and Sample Collection

Heart, blood, and tumor specimens were collected from cancer patients with irMyocarditis and ICI-treated controls without irMyocarditis (**Figure 1a-c, Methods**). Heart tissue for scRNA-seq was collected as part of clinically indicated endomyocardial biopsies in patients with concern for irMyocarditis (n = 15 biopsies; 13 with evidence of irMyocarditis and 2 negative biopsies used as controls) or as part of rapid autopsy collections (n = 4 autopsies; 3 with irMyocarditis and 1 control) (**Figure 1b, Supplementary Table 1-2**). Most endomyocardial biopsies were obtained from the right ventricular side of the interventricular septum as part of the clinical evaluation for suspected irMyocarditis, while autopsy samples were obtained by an autopsy pathologist from the right ventricular free wall (**Methods**). Of the irMyocarditis myocardial samples, 13 were collected before the initiation of corticosteroids; one patient (SIC_264) with fatal irMyocarditis provided one sample from a biopsy prior to the initiation of corticosteroids (denoted “SIC_264_A”) and another sample from autopsy after multiple lines of immunosuppression (“SIC_264_B”) (**Figure 1b**). The irMyocarditis patients in our heart scRNA-seq cohort were a median age of 72 years, 80% male, and represented eight primary tumor histologies (**Figure 1a**, **Supplementary Table 1**). All patients received a PD-1/PD-L1 inhibitor, of which 47% (n = 7) were treated with dual PD-1/CTLA-4 inhibition (**Supplementary Table 1**). Matched heart tissue, tumor, and normal tissues were collected, where possible, from irMyocarditis (n = 6) and ICI-treated control (n = 4) autopsy cases, and these were subsequently used for immunohistochemistry and bulk TCR-β sequencing experiments (**Figure 1a, 1c**, **Supplementary Table 3**). To evaluate correlations between tissue and blood, we captured both peripheral blood mononuclear cells (PBMCs) and serum for analysis **(Figure 1a, 1c**). PBMCs were collected from two ICI-treated control patients and 25 irMyocarditis patients: 13 patients included in our heart scRNA-seq dataset and 12 irMyocarditis patients from whom heart scRNA-seq data was not generated (**Supplementary Table 4**). In these patients, we collected samples prior to the start of corticosteroids for the treatment of irMyocarditis (“pre-corticosteroid”; n = 18 samples) and after the initiation of corticosteroids (“post-corticosteroid”; n = 26 samples) (**Supplementary Tables 5, 6**). Blood samples from prior to ICI treatment (“pre-ICI”; n = 7 samples) or after the initiation of ICI but before the emergence of clinical irMyocarditis (“on-ICI”; n = 4 samples) were available through archival biobanking efforts for a subset of cases and were subsequently retrieved **(Supplementary Table 6)**. In total, 55 PBMC specimens across timepoints were collected. Serum for secreted protein analysis was collected from irMyocarditis patients prior to the administration of corticosteroids (n = 16 patients) and ICI-treated control patients without irAEs (n = 10 patients) (**Supplementary Table 7**).

**Figure 1.**
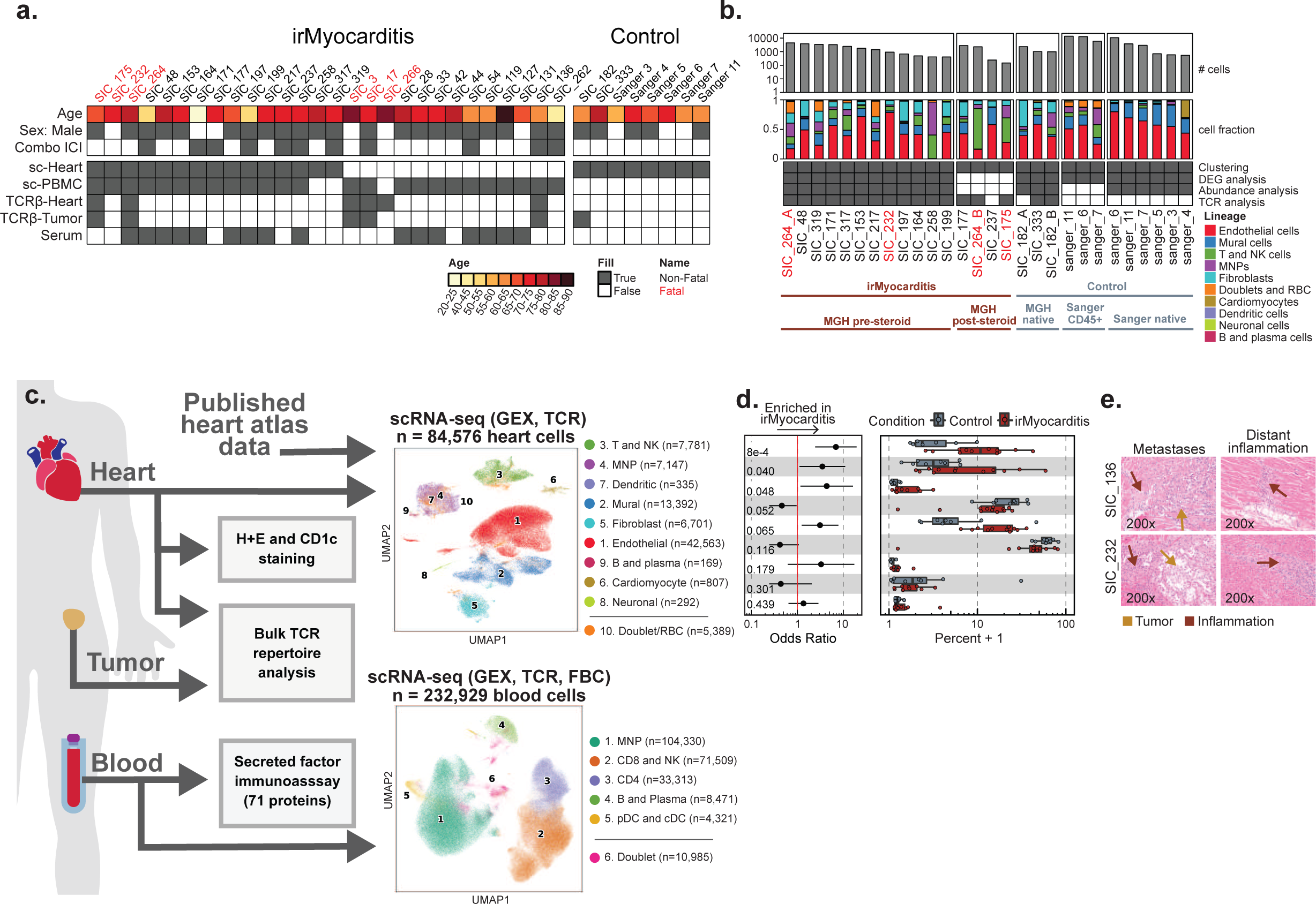
Multiple intracardiac cell populations are enriched in irMyocarditis. **a,** Patient cohort overview, including age, sex, ICI administered, and sample types obtained from irMyocarditis patients and controls. **b,** Top: bar plots displaying the number of single cells recovered from each heart sample. Middle: per donor color-coded cell lineage composition captured within each heart specimen with corresponding cell lineage color classifications indicated on the right. Bottom: grid depicting the contribution of each sample to the different analyses, grouped by sample characteristics and origins. Patients SIC_182 and SIC_264 each had two myocardial samples collected, the first collected at time of a diagnostic endomyocardial biopsy (labeled as sample A) and the second collected at time of autopsy (labeled as sample B). **c,** Left: summary of sample utilization across experimental frameworks. Right: UMAP embedding of scRNA-seq data from heart (top) and blood (bottom), color-coded by cell lineage, reporting number of cells per defined lineage. Cell lineage number was assigned according to the absolute number of cells detected per lineage and listed in order of ascending p-value for the associated analyses in panel **d** (for heart samples). **d,** Cell subset differential abundance analysis of major cell lineages comparing irMyocarditis cases (n = 12, red) to control in (n = 8, blue). Left: logistic regression odds-ratio for association of lineage abundance with irMyocarditis (OR > 1) vs control (OR < 1) for each lineage. Unadjusted likelihood-ratio test p-values for each lineage are shown. Right: boxplots show per patient intracardiac compositions of cell lineages where each dot represents a patient. Composition is reported as the percent of cells from a patient in each lineage. **e,** Hematoxylin and eosin (H&E) staining of cardiac tissues obtained via autopsy, SIC_136 (top, melanoma) and SIC_232 (bottom, renal cell carcinoma), showing intracardiac metastasis (left) and evidence of inflamed myocardium remote from metastatic foci (right). In **d,** error bars represent 95% confidence intervals. Boxes represent the median (line) and interquartile range (IQR) with whiskers extending to the remainder of the distribution, no more than 1.5x IQR, and each dot representing individual samples. Throughout the figure, red labels denote samples from patients with fatal irMyocarditis. Abbreviations: MNP, mononuclear phagocyte; GEX, gene expression; TCR, T cell receptor; FBC, feature bar code.

### scRNA-seq identifies increased T/NK, mononuclear phagocyte (MNP), and dendritic cells (DCs) in human irMyocarditis heart tissue

scRNA-seq analysis of our irMyocarditis heart samples yielded transcriptomes from 33,145 single cells from heart specimens. For additional control samples, we combined our data with six additional heart specimens (total 51,431 cells) from a published heart atlas that profiled donors without cancer and not receiving ICI therapy^29^ (**Figure 1c**). From this heart atlas, we utilized unenriched and CD45^+^ enriched samples from the right ventricular septum or right ventricular free wall to match the anatomic source of the majority of our irMyocarditis samples. Our integrated dataset included 84,576 single heart cells that passed quality control (QC) and was used for downstream analysis (**Methods**, **Supplementary Table 2**). We first employed iterative single-cell clustering strategies to define distinct cell populations. We subsequently leveraged differential abundance, differential gene expression, and cell-cell interaction analyses to identify cellular populations, cellular circuits, and transcriptional programs associated with irMyocarditis across cell lineages, which is further described below.

After data integration and unsupervised clustering analysis were performed at low resolution to identify all main cell lineages, the clustering analysis was repeated to define all cell subsets present within each lineage (**Methods)**. Nine broad cell lineages were identified in the heart specimens: T and NK cells (characterized by the expression of *CD3D* and *KLRB1*), MNPs (*CD14*, *CD68*), DCs (*CD1C*, *CLEC9A*, *IRF8*), B cells and plasmablasts (*CD79A*, *MZB1*), endothelial cells (*VWF, CA4*), mural cells (*RGS5*, *KCNJ8*), fibroblasts (*DCN*, *PDGFRA*), cardiomyocytes (*TNNI3*, *MB*), and neuronal cells (*PLP1*, *NRXN1*) **(Figure 1c-d, Supplementary Figure 1a, Supplementary Table 8**). Samples from our dataset and the published heart atlas^29^ contributed to each lineage (**Supplementary Figure 1b**) and to the 32 cell subsets defined through the subclustering of each cell lineage (**Supplementary Table 9**).

Given prior reports that corticosteroids can reverse the inflammatory processes of irAEs^30,31^ and decrease circulating inflammatory proteins in irAE patients^32^, our abundance analyses compared irMyocarditis samples collected prior to the administration of corticosteroids (“pre-corticosteroid” samples, n = 12) to control samples (n = 8) to define the cellular populations associated with disease. At the lineage level, this analysis showed an increase in the T/NK lineage (OR 6.8; 95% CI 2.4 - 19.2; p = 8.1e-4) and MNP lineage (OR 3.5; 95% CI 1.1 - 10.9; p = 0.040) in irMyocarditis cases (**Figure 1d, Supplementary Figure 1c, Supplementary Table 10**). Conventional dendritic cells (cDCs) were also more abundant in irMyocarditis (OR 4.3; 95% CI 1.2 - 16.1; p = 0.048). Trends towards increased fibroblasts (OR 3.1; 95% CI 1.2 - 7.8; p = 0.065) and decreased mural cells (OR 0.46; 95% CI 0.22 - 0.96; p = 0.052) in irMyocarditis samples were also observed. Similar to histologic reports of irMyocarditis^19^, we identified few intracardiac B cells/plasmablasts (n = 169 cells across cases and controls); since this lineage was rare and not enriched in irMyocarditis, they were not analyzed further.

### Microscopic tumor deposits are found in some patients with irMyocarditis

Beyond these shifts in cellular abundance associated with disease, we also found that two patients with irMyocarditis had evidence of viable tumor in the myocardium at autopsy, representing a potential driver of local inflammatory responses. One patient had a solitary lesion (SIC_136, melanoma), while the other had diffuse tumor deposits throughout the myocardial tissue (SIC_232, renal cell carcinoma); in both cases, there was inflammation both adjacent to and remote from the metastases (**Figure 1e**). A third patient with metastatic melanoma and irMyocarditis who had received prior targeted therapy before dual checkpoint inhibition (SIC_171) had evidence of melanin-laden macrophages on endomyocardial biopsy, suggesting prior intracardiac melanoma (*data not shown*). While microscopic metastases in the setting of a suspected irAE have been described in the gut^33^, intracardiac tumor deposits in the vicinity of irMyocarditis previously have not been reported. While we did not find evidence of viable tumor cells in our scRNA-seq data, these histological findings suggest that inflammation against intracardiac tumors may contribute to myocardial inflammation in a subset of patients who present with irMyocarditis.

### scRNA-seq of blood identifies circulating immune cell populations that correlate with irMyocarditis severity

PBMCs were profiled using scRNA-seq, paired TCR, and surface proteomics using the CITE-seq protocol^34^ for 197 surface proteins (**Supplementary Table 11**) to identify peripheral blood correlates of irMyocarditis and disease severity as well as to examine the relationship between irAE tissue and blood. Blood samples were obtained over multiple time points from 25 patients (n = 55 total samples) who developed irMyocarditis and two ICI-treated control patients who never developed irMyocarditis (**Supplementary Figure 1d, Supplementary Table 4**)^35^. Unsupervised clustering of 232,929 cells that passed QC filters (**Methods**) identified all known blood immune cell lineages, defined by canonical markers: CD8 T cells and NK cells (*CD3D*, *CD8A*, *KLRF1*); CD4 T cells (*CD3D, CD4*); MNPs (*LYZ*, *CD14)*; DCs, representing both plasmacytoid dendritic cells (pDCs) (*LILRA4*) and cDCs (*CD1C*, *CLEC9A*); and B and plasmablast cells (*CD79A*, *JCHAIN*) (**Figure 1c, Supplementary Figure 1e, Supplementary Tables 6, 9, and 12**).

We then sought to identify populations in the blood associated with irMyocarditis disease severity by investigating the relationship between the circulating frequency of each lineage and serum troponin levels, a widely used clinical measure that is associated with poor outcomes in irMyocarditis^5,36^. The peripheral abundance of MNPs was directly correlated with serum troponin (p = 0.001), while CD4 T cells (p = 0.024) and CD8 T/NK cells (p = 0.003), and B/plasmablast cells (p = 0.039) were inversely correlated (**Supplementary Figure 1f, Supplementary Table 16**). These findings are consistent with previous reports of lymphopenia in the setting of irMyocarditis^37^ but provide greater granularity to nominate novel peripheral blood correlates of irAE disease severity.

### Activated CD8 T cells and CD4 T cells are increased in irMyocarditis

In order to further understand the nature of the T cells present in irMyocarditis heart tissue, we subclustered the 9,134 cells in the intracardiac T/NK lineage. We annotated cell subtypes by using marker genes (i.e., area under the curve [AUC] ≥ 0.75, one-vs-all [OVA] pseudobulk false discovery rate [FDR] < 0.05; **Methods, Supplementary Table 8**) and cross-referencing marker genes with published immune transcriptional signatures^29,38,39^. Throughout the text, immune cell subsets found in heart tissue are denoted by subset names that begin with “h-”, while subsets in blood begin with “b-”. Within the T/NK lineage, we defined six distinct cell subsets, including four CD8 T, one CD4 T (cluster 3, h-CD4T*^IL7R^*^,*LTB*^), and one NK cell subset (cluster 1, h-NK*^KLRF1,FCER1G^*) (**Figure 2a-c, Supplementary Table 9**). All CD8 T cell subsets broadly expressed markers of cytotoxicity (*e.g., GZMK* and *CCL5*) and cell adhesion (e.g., *ITGB7*) (**Figure 2d)**. Two CD8 T cell subsets – cluster 2 (h-CD8T*^CD27,LAG3^),* which is defined by the expression of *CD27* and *LAG3*, and cluster 6 (h-*CD8T*^cycling^), which is defined by a cycling cell signature (*e.g.*, *STMN1*, *TOP2A*) – expressed a spectrum of activation and exhaustion markers (*e.g., CTLA4, PDCD1, and TOX*) as well as the chemokine receptor *CXCR3*. Cluster 4 (h-CD8T*^CCL5,NKG7^*) captured a CD8 T cell population expressing *CCL5* and *NKG7* and also included a small group of cycling cells with lower expression of exhaustion markers like *LAG3* and *TOX*. Finally, cluster 5 (h-CD8T*^KLRG1,CX3CR1^*) comprises CD8 T cells distinctly expressing *KLRG1* and *CX3CR1,* which are markers known to define short-lived effector cells^40^. Amongst these T cell subsets, h-CD8T*^CD27,LAG3^* (OR 5.8; 95% CI 1.3 - 26.3; p = 0.03), h-CD8T*^CCL5,NKG7^* (OR 19.0; 95% CI 6.0 - 60.0; p = 5.1e-05), and h-CD4T*^IL7R^*^,*LTB*^ (OR 6.0; 95% CI: 1.8 - 20.1; *p* = 0.0091) were more abundant in irMyocarditis cases than controls, while h-CD8T^cycling^ displayed a trend towards greater abundance (OR 11.7; 95% CI 1.3 - 104.0; p = 0.05) (**Supplementary Figure 2a-b, Supplementary Table 10**). Although our scRNA-seq analyses of the heart were underpowered to compare fatal and non-fatal cases, we used serum troponin as a marker of disease severity^5,36^ and found that the intracardiac frequency of the h-CD8T^cycling^ subset correlated with serum troponin (p = 0.018) (**Figure 2e, Supplementary Table 17**). Marker genes that defined h-CD4T*^IL7R^*^,*LTB*^ include a mixture of naive (*CCR7* and *MAL*), regulatory (*IL2RA* and *FOXP3*), and memory (*LTB* and *FLT3LG*) CD4 T cells^39^, but given the small number of cells in the h-CD4T*^IL7R^*^,*LTB*^ subset (n = 1,401 cells), further subsetting analysis was not explored.

**Figure 2.**
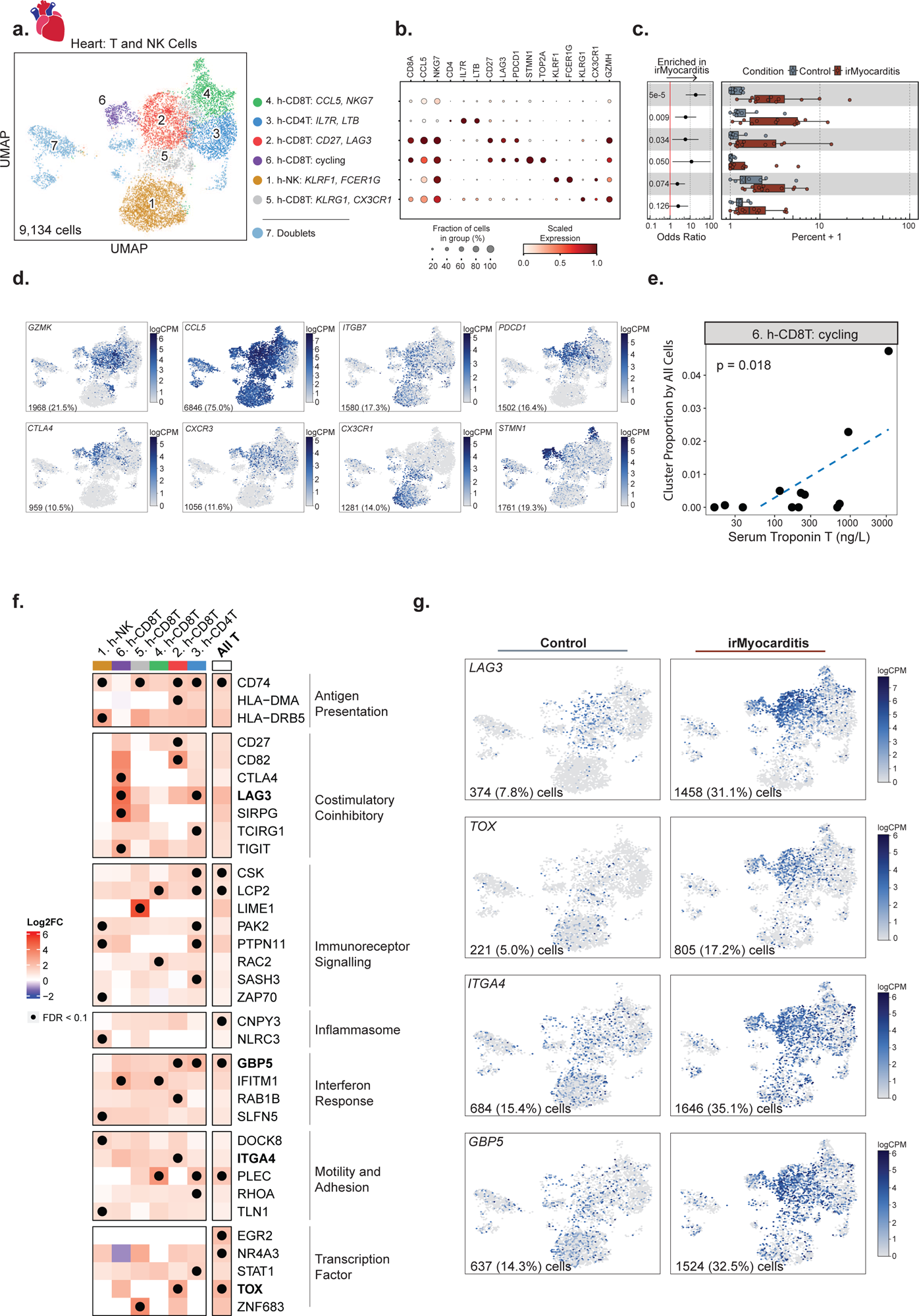
Activated, cytotoxic T cells are enriched in irMyocarditis heart tissue. **a,** UMAP embedding of 9,134 CD8 T, CD4 T, and NK cells recovered from scRNA-seq of heart tissue, colored by the seven defined subsets labeled on the right. Cell subset number was assigned according to the absolute number of cells detected per subset and are listed in order of ascending p-value for the associated analyses in panel **c**. **b,** Dot plot showing selected marker genes for each T and NK cell subset. Dot size represents the percent of cells in the subset with non-zero expression of a given gene. Color indicates scaled expression. **c,** Cell subset differential abundance analysis comparing irMyocarditis cases (n = 12, red) to control (n = 8, blue). Left: logistic regression odds-ratio for association of cell subset abundance with irMyocarditis (OR > 1) versus control (OR < 1) for each subset. Unadjusted likelihood-ratio test p-values for each cell subset are shown. Right: boxplots show per patient intracardiac compositions of T/NK cell subsets, where each dot represents a patient. Composition is reported as the percent of T/NK cells from a patient in each subset. **d,** Feature plots using color to indicate gene expression (logCPM) levels of the indicated genes projected onto the T/NK UMAP embedding. Cell numbers and percentages represent gene expression across all heart T and NK cells. **e,** h-*CD8T*^cycling^ subset intracardiac abundance (y-axis) versus serum troponin T (x-axis) for irMyocarditis patients (n = 12). Linear regression p value shown. **f,** Heatmap showing selected differentially expressed genes (irMyocarditis versus control) across the T/NK subsets grouped by biological categories. The “All T” row depicts differential gene expression results from a pseudobulk analysis of pooled T-cell subsets (subsets 2-6) and excludes NK cells. Color scale indicates log2 fold-change difference between irMyocarditis cases and controls. Black dots indicate FDR < 0.1 (Wald test). Genes indicated in bold are displayed in panel **g**. **g,** Feature plots using color to indicate gene expression (logCPM) levels of the indicated genes expressed by control (left) or irMyocarditis samples (right), projected onto the T/NK UMAP embedding. Cell numbers and percentages represent gene expression across T/NK cells in control or irMyocarditis samples. In **c**, error bars represent 95% confidence intervals; boxes represent the median (line) and interquartile range (IQR) with whiskers extending to the remainder of the distribution, no more than 1.5x IQ, and dots represent individual samples.

### Immunosuppression alters T-cell gene expression but not TCR repertoire in an irMyocarditis case with matched pre- and post-corticosteroid heart samples

While immunosuppressive agents such as corticosteroids and abatacept are used to treat irMyocarditis^15,30,31,41^, there is no evidence as to how these agents impact immune cell populations or transcriptional programs that persist despite immunosuppressive treatments. Clinical opportunities to investigate paired pre-corticosteroid and post-corticosteroid heart samples are rare, but they could help to shape the interpretation of data collected from post-corticosteroid samples in our study and irAE efforts that similarly leverage autopsy samples^19,23^. We analyzed a single patient (SIC_264) with fatal irMyocarditis who provided both a pre-corticosteroid biopsy specimen (SIC_264_A) and a second sample (SIC_264_B) obtained at autopsy after the administration of high dose corticosteroids (a total of 6 g methylprednisolone over seven days) and abatacept (10 mg/kg administered two days prior to autopsy) (**Figure 1b**). Across these two samples, the top 13 most expanded TCR clones were found in both timepoints, suggesting that the most predominant TCR-β sequences remain similar before and after immunosuppressive treatment in the setting of persistent irMyocarditis (**Supplementary Figure 2c-d**). However, gene expression profiles of the cells carrying these TCR-β sequences appear to change post-corticosteroid treatment. Prior to the initiation of corticosteroids, these expanded TCR-β clones were predominantly mapped to CD8 T cells expressing cycling markers (e.g. *STMN1*) in the h-*CD8T*^cycling^ and h-CD8T*^CCL5,NKG7^* subsets (**Supplementary Figure 2d-e**). At the time of death, seven days later, these expanded TCR-β clones were chiefly mapped to h-CD8T*^CD27,LAG3^*, which express activation markers but do not express a cycling signature (**Figure 2d**, **Supplementary Figure 2d-e**). These observations suggest the possibility that immunosuppression alters the cellular phenotype and transcriptional profile of T cells even if they do not dramatically alter the predominant TCR clones in the setting of treatment-resistant irMyocarditis. This observation provided further justification for utilizing only pre-corticosteroid samples for our differential gene expression and abundance analyses. Nonetheless, given that no notable changes post-corticosteroid were observed across the top expanded TCR clones, the post-corticosteroid samples were kept in downstream TCR repertoire analyses.

### T cells in irMyocarditis show transcriptional signatures consistent with increased immunoreceptor signaling, interferon response, and motility

To identify transcriptional programs present in intracardiac T and NK cells, we performed differential gene expression analysis comparing irMyocarditis cases versus controls at the level of T cell lineage (“All T” cells) and for each of the defined subsets. As T cells in the uninflamed heart are rare^29,41^, we included in the analysis ICI-treated controls without irMyocarditis from MGH (n = 2), as well as the native and CD45^+^ enriched samples from non-ICI-treated controls (n = 6)^29^. Significant upregulation (FDR<0.1) of transcriptional programs associated with MHC class II presentation (*CD74, HLA-DMA, HLA-DRB5*), immunoreceptor signaling (*LCP2, LIME1, PAK2, NR4A3, EGR2, TOX*), and interferon responses (*GBP5, IFITM1, STAT1*) were seen across multiple subsets (**Figure 2f-g, Supplementary Table 14**). Additionally, inflammasome genes (*CNPY3*, *NLRC3*) were upregulated across all T cell subsets and NK cells. In the h-CD8T*^CD27,LAG3^* cells, upregulation of multiple immune checkpoints (*CTLA4* and *LAG3*) were observed. Other genes involved in cell migration and adhesion, such as *ITGA4* in the h-CD8T*^CD27,LAG3^*subset, were also increased in irMyocarditis cases.

### TCR clones enriched in irMyocarditis hearts are distinct from those enriched in tumor

We next sought to assess TCR clonality in irMyocarditis myocardial tissue and to characterize their relationship to TCR clones present in tumor tissue from the same patients after ICI treatment. Bulk TCR-β sequencing^42^ allows for deeper sequencing of the TCR-β repertoire than single-cell (sc)TCR-seq (**Figure 3a**, **Supplementary Table 3, Methods**). Of the six patients without diffuse myocardial metastases, the heart tissues were classified by a cardiac pathologist as “active myocarditis” (n = 2; SIC_17 and SIC_264), lymphocytic infiltrate without associated myocyte injury suggestive of irMyocarditis (“borderline”; n = 1; SIC_136), “healing myocarditis” (n = 3; SIC_3, SIC_175, and SIC_266), or no evidence of myocarditis/control (n = 4) according to the Dallas Criteria^43^. Areas of interest were dissected on serial FFPE sections, and gDNA was extracted for bulk TCR-β sequencing (**Methods**). As expected, irMyocarditis tissues yielded more TCR-β sequences per total nucleated cells (control mean 0.9%±0.6 SD vs irMyocarditis mean 17.7%±14.5 SD; p = 0.00023) (**Supplementary Figure 3a**). To compare the TCR-β diversity of our irMyocarditis tissue repertoires, we used Hill’s diversity index across different diversity orders to capture both the true richness (low q value) and abundance (high q value) of the samples (**Figure 3b**). Notably, the two “active” irMyocarditis cases had the lowest diversity across all orders, and the TCR-β repertoire from these two patients each demonstrated expansion of primary dominant clones that comprised >40% of the recovered irMyocarditis TCR-β repertoire (**Supplementary Figure 3b-c**).

**Figure 3.**
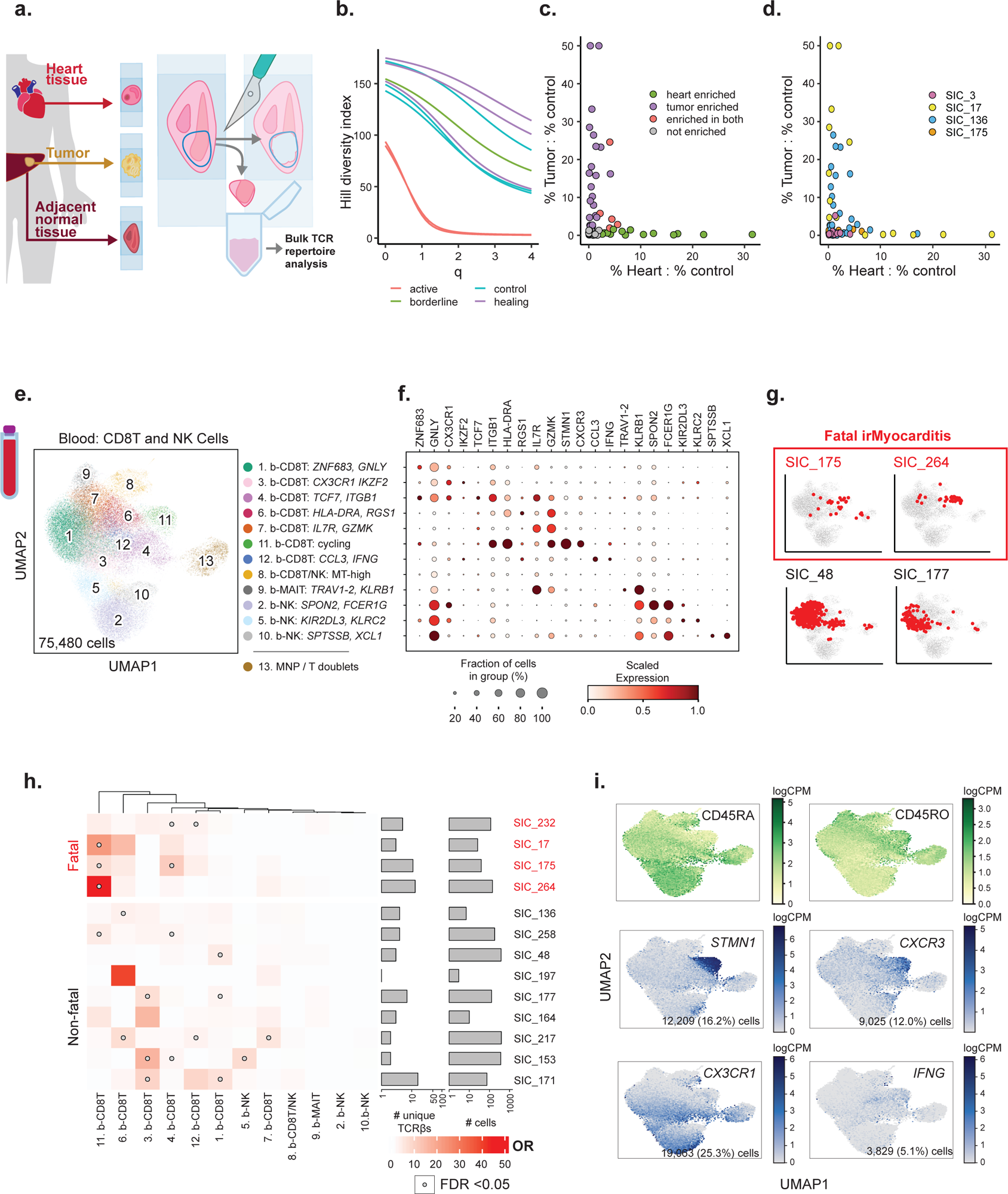
T-cell receptors (TCR) distinguish anti-tumor responses from irMyocarditis and identify markers of fatal irMyocarditis in blood. **a,** Schematic of the TCR-β chain sequencing experiment. Paired irMyocarditis, ICI-treated tumor, and/or normal parenchyma adjacent to tumor (“control tissue”) was marked and macroscopically dissected from serial slides. Excised portions were processed and underwent TCR-β chain sequencing. **b,** Smoothed Hill’s diversity index curves at diversity orders 0-4 for the TCR-β repertoires of irMyocarditis tissues, colored by histologic appearance at the time of autopsy. **c, d,** The proportion of each expanded TCR-β clone in heart and tumor tissue was calculated and then normalized by dividing by the proportion of the same clone in control tissue. Normalized TCR-β clone proportions for heart (x-axis) and tumor tissue (y-axis) are plotted. Individual data points considered significant (FDR < 0.05, Fishers exact test) are colored by tissue(s) of enrichment (**c**) and by donor (**d**). **e**, UMAP embedding of 75,480 CD8 T and NK cells recovered from scRNA-seq of peripheral blood cells, colored by the 13 defined cell subsets labeled on the right. Subset number was assigned according to the absolute number of cells detected per subset and ordered by abundance within major cell types. **f,** Dot plot showing selected marker genes for each CD8 T and NK cell subsets defined in blood. Dot size represents the percent of cells in the subset with non-zero expression of a given gene. Color indicates scaled expression across subsets. **g**, TCR-β sequences were compared between blood and heart T cells. Cells in blood for which the same TCR-β was expanded in the irMyocarditis heart sample from the same patient are shown in red on CD8 T/NK blood UMAP embeddings from representative patients. **h,** Logistic regression was used to determine associations between blood CD8 T cell subsets and the presence of expanded TCR-β clones from irMyocarditis heart samples in those subsets. Far left: heatmap showing the calculated associations, displayed as odds ratios. Color in the heatmap represents the magnitude of the regression odds ratio. Open circles indicate FDR < 0.05 (likelihood ratio test). Middle: the number of unique heart-expanded TCR-β sequences found to be shared with blood. Right: total number of cells containing those TCR-β sequences for each donor. **i**, Top: feature plots using color to indicate surface protein levels (logCPM) of CD45RA and CD45RO protein (as determined by CITE-seq) projected onto the blood CD8 T and NK cell UMAP embedding. Bottom: feature plots using color to indicate gene expression (logCPM) levels of the indicated genes projected onto the blood CD8 T and NK cell UMAP embedding. Cell numbers and percentages represent gene expression across all blood CD8 T and NK cells. Throughout the figure, red labels denote samples from patients with fatal irMyocarditis.

As prior reports have suggested TCR sharing between irMyocarditis heart tissue and autologous tumor^19,44^, we compared the TCR-β repertoires in irMyocarditis heart to autologous tumors. To account for ubiquitous, circulating “bystander” T cells^45^ and resident T-cell populations in normal parenchyma that had been invaded by tumors^46^, we also analyzed the TCR-β repertoire of normal tissue adjacent to tumors (available from four donors) to establish a common control tissue against which the heart and tumor tissue could be compared (**Methods**). We considered TCR-βs to be enriched in the tumor or heart if they were expanded in either of those tissues relative to the TCR-β frequency in the matched normal control tissue. Heart-enriched TCR-β clones were recovered from all four donors with irMyocarditis (range 1-9 per donor; total 19 clones), while tumor-enriched clones were found in three of these four donors (range 0-17; total 28 clones). Across all irMyocarditis patients, five clones were enriched in both heart and tumor (18% of tumor-enriched TCR-βs, 26% of heart-enriched TCR-βs), but in each irMyocarditis patients, the TCR-β clones most expanded in heart tissue compared to control tissue were not enriched in the tumor relative to control tissue (**Figure 3c-d, Supplementary Figure 3d-e, Supplementary table 18**). In sum, these data suggest that there exists a modest overlap of TCR-β clones between heart and tumor, but the majority of the enriched TCR-β clones and the most expanded clones in each tissue are distinct.

### Fatal irMyocarditis is associated with cycling CD8 T cells in the blood that express TCRs expanded in intracardiac T cells

We next examined if TCR-β clones enriched in irMyocarditis heart tissue could also be detected among circulating CD8 T cells, which would suggest that intracardiac T cells may have emigrated from clonally related populations in blood. We subclustered 75,480 CD8 T/NK cells from our peripheral blood dataset and identified 12 distinct cell subsets (**Figure 3e-f, Supplementary Figure 4a, Supplementary Table 9**). We combined our heart single-cell data with TCR-β sequence data yielded from the analysis of bulk tissue (**Figure 3a**) from patients with “active” (SIC_17, SIC_264) or “borderline” irMyocarditis (SIC_136) (**Supplementary Figure 3b**). Across all patients, logistic regression analysis showed statistically significant sharing of TCR-β sequences between the heart and five blood CD8 T subsets (**Supplementary Figure 4b-c**). However, there were striking differences in heart/blood TCR sharing seen across specific CD8 T cell subsets on a per-patient basis (**Figure 3g-h, Supplementary Figure 4d**). Three of four fatal irMyocarditis patients had statistically significant sharing of heart-expanded TCR-β clones in the circulating b-CD8T^cycling^ cells while only one of nine patients with non-fatal irMyocarditis had significant sharing in this subset (p = 0.052 for this comparison by Fisher’s exact test). b-CD8T^cycling^ is defined by expression of *CXCR3*, cycling genes (*MKI67, STMN1*), and CD45RO surface protein, indicating that these are antigen-exposed and dividing cells (**Figure 3i**). The only patient with fatal irMyocarditis who did not share clones between the heart and the circulating b-CD8T^cycling^ subset was a patient with diffuse tumor deposits in the heart (SIC_232). We then sought to identify where the expanded heart TCR-β clones mapped across the intracardiac T-cell subsets; of the cases with >10 intracardiac T cells sharing clones between heart and blood, shared TCRs predominantly mapped to *CXCR3*-expressing subsets in a fatal case (SIC_264) and to *CX3CR1*-expressing subsets in three non-fatal cases (**Supplementary Figure 4e-f**). These results suggest that the phenotypes of T cells found shared between the heart and blood may risk-stratify patients with irMyocarditis and shed light on circulating blood CD8 T cells that may contribute to disease pathogenesis.

We then sought to perform similar analysis in blood CD4 T cells. We subclustered 33,313 blood CD4 T cells and identified six distinct cell subsets (**Supplementary Figure 5a-c, Supplementary Tables 9, 13).** In contrast to circulating CD8 T cells, there was minimal TCR-β sharing between blood and intracardiac CD4 T cells. No patient had more than ten blood CD4 T cells sharing a TCR-β with an expanded intracardiac T cell clone (*data not shown*).

### cDC enrichment in irMyocarditis hearts is associated with disease severity

We next investigated intracardiac mononuclear phagocyte (MNP) populations. Unsupervised clustering of 9,824 cells yielded seven unique populations: five distinct MNP subsets, conventional dendritic cells (cDCs), and plasmacytoid dendritic cells (pDCs) (**Figure 4a-c, Supplementary Table 9**). Expression of *LYVE1* and *C1QA* distinguished one MNP subset (cluster 2), which likely represents cardiac resident macrophages^47,48^. MNP cluster 3 expressed *FCGR3A*, and the immune checkpoint molecules *LILRB1* and *LILRB2*, which confer inhibitory signals to MNPs (**Figure 4b, 4d**)^49,50^. Additional MNP populations were characterized by the expression of *S100A8*, *S100A12*, and *VCAN* (cluster 4), and the expression of *TREM2* and *APOC1* (cluster 5) (**Figure 4b**). Among cDCs (cluster 6, defined by *CLEC9A* and *CD1C* expression), a subset expressed *FLT3,* which is a critical homeostatic signal for DC maintenance^51^ (**Figure 4d**). There were too few h-cDCs^CLEC9A,CD1C^ detected to further subcluster these cells into cDC1 (*CLEC9A*) and cDC2 (*CD1C*), though we noted that the majority of h-cDCs ^CLEC9A,CD1C^ observed expressed *CD1C* (**Figure 4d**). Multiple chemokine receptors were expressed across intracardiac MNP populations. The chemokine receptor *CXCR4*, which controls DC activation and localization^52^, was highly expressed by h-cDCs^CLEC9A,CD1C^, h-pDCs^LILRA4, IRF8^, and a fraction of h-MNP^FCGR3A,LILRB2^, h-MNP^S100A12,VCAN^, and h-MNP^TREM2,APOC1^. CCR2, which supports monocyte recruitment early after cardiac injury ^53–56^, was expressed only by a small fraction of h-MNP^FCGR3A,LILRB2^ and h-MNP^S100A12,VCAN^. *CX3CR1*, which has been shown to attenuate the severity of viral myocarditis ^57^ and is expressed by murine cardiac macrophages ^54,58^, was lowly expressed in a subset of the h-MNP^FCGR3A,LILRB2^ population. Two MNP subsets, h-MNP^S100A8-low,C1QA-low^ (cluster 1; OR 4.9; 95% CI 1.3 - 18.1; p = 0.044) and h-MNP^FCGR3A,LILRB2^ (cluster 3; OR 3.3; 95% CI 1.2 - 9.3; p = 0.041), and h-cDCs ^CLEC9A,CD1C^ (OR 3.5; 95% CI 1.1 - 11.9; p = 0.064) were more abundant in irMyocarditis versus control hearts (**Figure 4c, Supplementary Figure 6a-b, Supplementary Table 10**). We recovered only 43 pDCs (cluster 7, defined by LILRA4 and IRF8), and while h-pDCs^LILRA4,IRF8^ may be more abundant in irMyocarditis cases (cluster 7; OR 4.34 x 10^7^; CI 97.46-1.93 x 10^13^; p = 0.018), such a low recovery does not allow us to confidently assess these differences.

**Figure 4.**
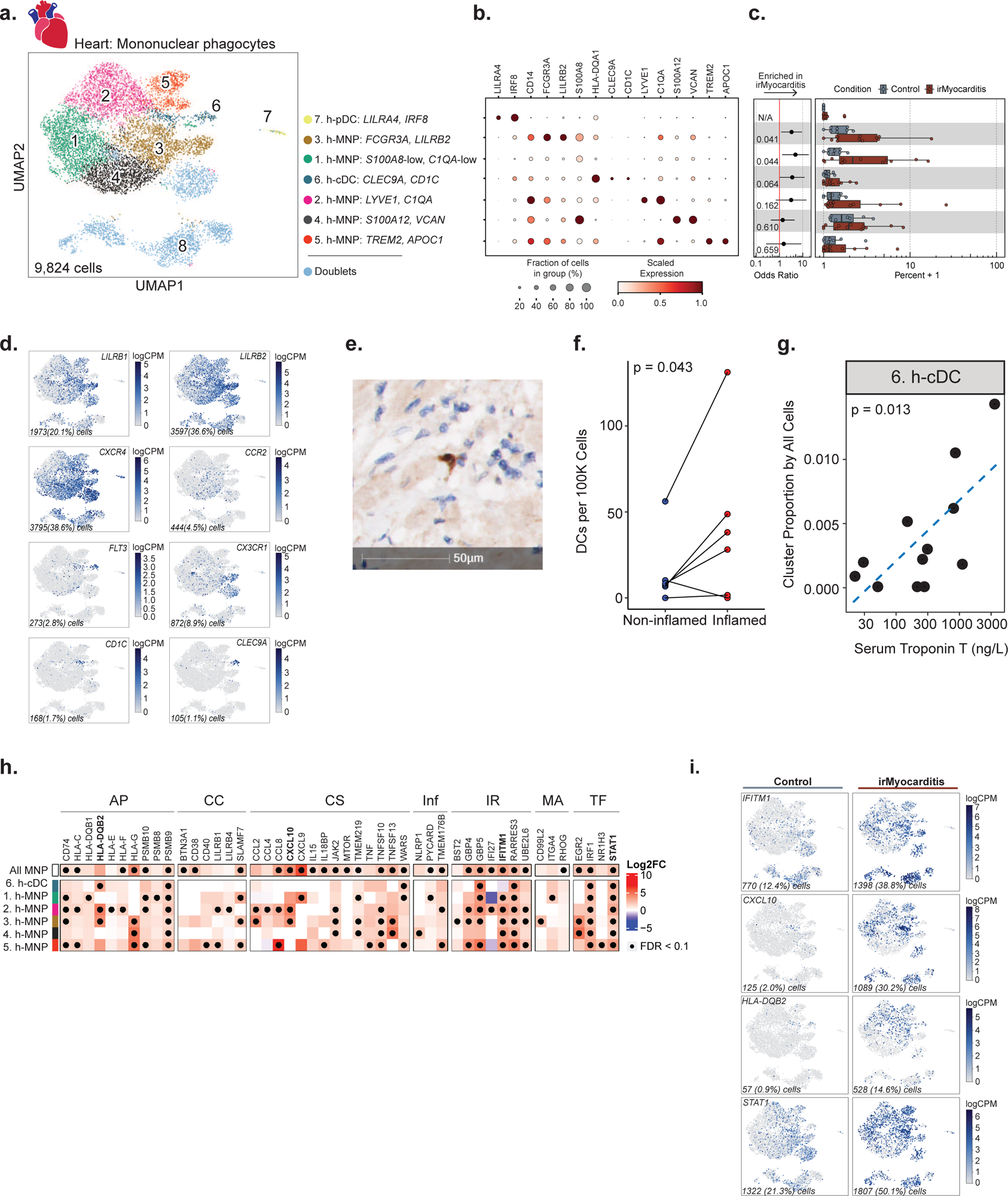
Conventional dendritic cells (cDCs) are enriched in irMyocarditis heart tissue and associated with disease severity. **a,** UMAP embedding of 9,824 mononuclear phagocytes (MNPs) isolated from heart tissue, colored by the eight defined cell subsets labeled on the right. Subset number was assigned according to the absolute number of cells detected per subset and are listed in order of ascending p-value for the associated analysis in panel **c**. **b**, Dot plot showing top marker genes for each MNP subset. Dot size represents the percent of cells in the subset with non-zero expression of a given gene. Color indicates scaled expression across subsets. **c,** Cell subset differential abundance analysis comparing irMyocarditis cases (n = 12, red) to control (n = 8, blue). Left: logistic regression odds-ratio for association of cell subset abundance with irMyocarditis (OR > 1) vs control (OR < 1) for each subset. Unadjusted likelihood-ratio test p-values for each subset are shown. Statistics for h-pDC^LILRA4, IRF8^ are not included due to extremely low recovery from this cell subset. Right: boxplots show per patient intracardiac MNP cell subset compositions for each subset, where each dot represents a patient. Composition is reported as the percent of cells from a patient in each subset. **d**, Feature plots using color to indicate gene expression (logCPM) levels of the indicated genes projected onto the heart MNP UMAP embedding. Cell numbers and percentages represent gene expression across all heart MNP cells. **e**, Representative image following immunohistochemical staining of irMyocarditis heart tissue section for CD1c (brown) and hematoxylin nuclear counterstaining (blue). **f**, Comparison of CD1c^+^ cell density as measured by immunohistochemical staining in non-inflamed regions (left column) and inflamed regions (right column) of irMyocarditis heart sections; p = 0.043, one-sided T-test. **g**, h-cDCs^CLEC9A,CD1C^ proportion (y-axis) versus serum troponin T (x-axis) for irMyocarditis samples (n = 12); p = 0.013, linear regression. **h**, Heatmap showing select differentially expressed genes (irMyocarditis versus control) in heart tissue grouped by the following biological themes: antigen presentation (AP), co-inhibition or co-stimulation (CC), cytokine signaling (CS), inflammasome (Inf), interferon response (IR), motility and adhesion (MA), and transcription factors (TF). The “All MNP” row depicts differential gene expression results from a pseudobulk analysis of pooled MNP subsets 1 through 5, excluding cDCs. Color scale indicates log2 fold-change difference between irMyocarditis cases and controls. Black dots indicate FDR < 0.1 (Wald test). **i**, Feature plots using color to indicate gene expression (logCPM) levels of the indicated genes expressed by control samples (left) or irMyocarditis samples (right) projected onto the heart MNP UMAP embedding. Cell numbers and percentages represent gene expression across respective control or irMyocarditis sample MNP cells. In **c**, error bars represent 95% confidence intervals. Boxes represent the median (line) and interquartile range (IQR) with whiskers extending to the remainder of the distribution, no more than 1.5x IQR, with dots representing individual samples.

We then sought to validate the increased abundance of cDCs in irMyocarditis heart tissue. Using autopsy samples, histologic sections of ICI-treated control heart tissue and irMyocarditis heart tissue were stained for the cDC marker CD1c, and CD1c^+^ cells were quantified using digital imaging software to highlight stained candidate cDCs, followed by manual validation (**Figure 4e, Methods**). Across whole slides, irMyocarditis tissue showed a trend towards increased CD1c^+^ cell density compared to control tissue (22.9±35.0 vs 2.5±2.1 CD1c^+^ cells per 10^5^ total cells; p = 0.11) (**Supplementary Figure 6c, Supplementary Table 19**). However, areas of inflammation in irMyocarditis hearts had more than 16-fold greater density of CD1c^+^ cells than control tissue (41.3±48.2 vs 2.5±2.1; p = 0.053) (**Supplementary Figure 6d**) and a 2.8-fold greater density of CD1c^+^ cells than non-inflamed regions of irMyocarditis hearts (41.3±48.2 vs 14.9±20.5; p = 0.043) (**Figure 4f**). Collectively, these IHC data support the finding from scRNA-seq data that cDCs are increased in areas of inflammation in irMyocarditis.

Given the observation of increased abundance of cDCs in irMyocarditis heart tissue, we investigated whether the frequencies of intracardiac MNP subsets were associated with irMyocarditis severity as measured by serum troponin. While h-cDCs^CLEC9A,CD1C^ frequency was found to be correlated with serum troponin (p = 0.013) (**Figure 4g**), similar relationships were not seen with the two other MNP populations enriched in irMyocarditis tissue (i.e., h-MNP^S100A8-low, C1QA-low^ and h-MNP^FCGR3A, LILRB2^; **Supplementary Table 17**).

### MNPs in irMyocarditis show transcriptional changes involved in antigen presentation, T cell activation, and immune cell recruitment

Beyond a shift in population abundance, several transcriptional programs were upregulated in irMyocarditis versus control tissue across multiple MNP subsets (**Figure 4h-i, Supplementary Table 14**). This included genes related to antigen presentation, such as MHC class I (*HLA-C*) and MHC class II (*HLA-DQB1, HLA-DQB2*) programs, as well as the proteasome subunit beta type-9 (*PSMB9*), an interferon-inducible component of the immunoproteasome^59^. Additionally, the immune checkpoint molecule *HLA-G* was upregulated in several MNP subsets^60^. Genes encoding CXCR3 ligands *CXCL9* and *CXCL10* were upregulated in a subset of MNPs and at the pseudobulk level across all MNP subsets. Furthermore, all MNP subsets and h-cDCs^CLEC9A,CD1C^ showed increased expression levels of interferon stimulated genes (ISGs). For example, consistent with a prior report^21^, the interferon-inducible inflammasome activator *GBP5* was increased in irMyocarditis, along with ISGs such as *IFITM1* and *RARRES3*. Finally, irMyocarditis samples demonstrated increased expression of the key mediator of cytokine signaling and transcription factor *STAT1.* Collectively, multiple transcriptional programs related to T cell activation and interferon responses were upregulated in the setting of irMyocarditis.

### Fibroblasts are enriched in irMyocarditis heart tissue

Given stromal cells such as endothelial cells, pericytes, and fibroblasts play crucial roles in orchestrating cardiac immune responses^61,62^, we next investigated the contribution of the non-immune cell component captured in our heart data to irMyocarditis. Subclustering of scRNA-seq data from 65,409 cells yielded 17 distinct non-immune subsets (excluding doublets and RBCs): eight endothelial populations, three pericyte populations, fibroblasts, myofibroblasts, smooth muscle cells, endocardial cells, neuronal cells, and cardiomyocytes (**Figure 5a-b, Supplementary Figure 7a-b, Supplementary Table 9, Methods**). Within the endothelial cell compartment, there were three capillary endothelial cell populations distinguished by *CA4* expression^63,64^, two arterial endothelial cell populations characterized by *HEY1* expression^65^, a population of inflammatory endothelial cells expressing multiple ISGs, such as *GBP5* and *IFITM1*, a venous endothelial population expressing marker *ACKR1*^66^, and lymphatic endothelial cells distinguished by *PDPN* and *CCL21* expression^63,67^. Amongst all non-immune subsets, only fibroblasts (defined by expression of *DCN* and *LUM*) were more enriched in irMyocarditis tissue (cluster 4; OR 3.8; 95% CI 1.5 – 9.7; p = 0.042) (**Figure 5c, Supplementary Figure 7c-d, Supplementary Table 10**). Multiple subsets were less abundant in irMyocarditis: Capillary EC*^RGCC,CA4^* (cluster 1; OR 0.39; 95% CI 0.23 – 0.69; p = 0.0061), Capillary EC*^CXCL2,JUN^* (cluster 2; OR 0.47; 95% CI 0.23 – 0.98; p = 0.063), Arterial EC^HEY1,SEMA3G^ (cluster 5; OR 0.43; 95% CI 0.21 – 0.88; p =0.037), Arterial EC^RBP7,BTNL9^ (cluster 9; OR 0.29; 95% CI 0.13 – 0.64; p = 0.0064), smooth muscle cells (cluster 8; OR 0.38, 95% CI 0.20 – 0.75; p = 0.0097), and Pericytes^KCNJ8,ABCC9^ (cluster 7; OR 0.50; 95% CI 0.27 – 0.91; p = 0.038). In sum, fibroblasts were uniquely enriched in irMyocarditis among all non-immune populations. The decrease in relative frequency of other non-immune populations (among all total measured intracardiac cells) may be due to the infiltration of immune cells in the setting of irMyocarditis.

**Figure 5.**
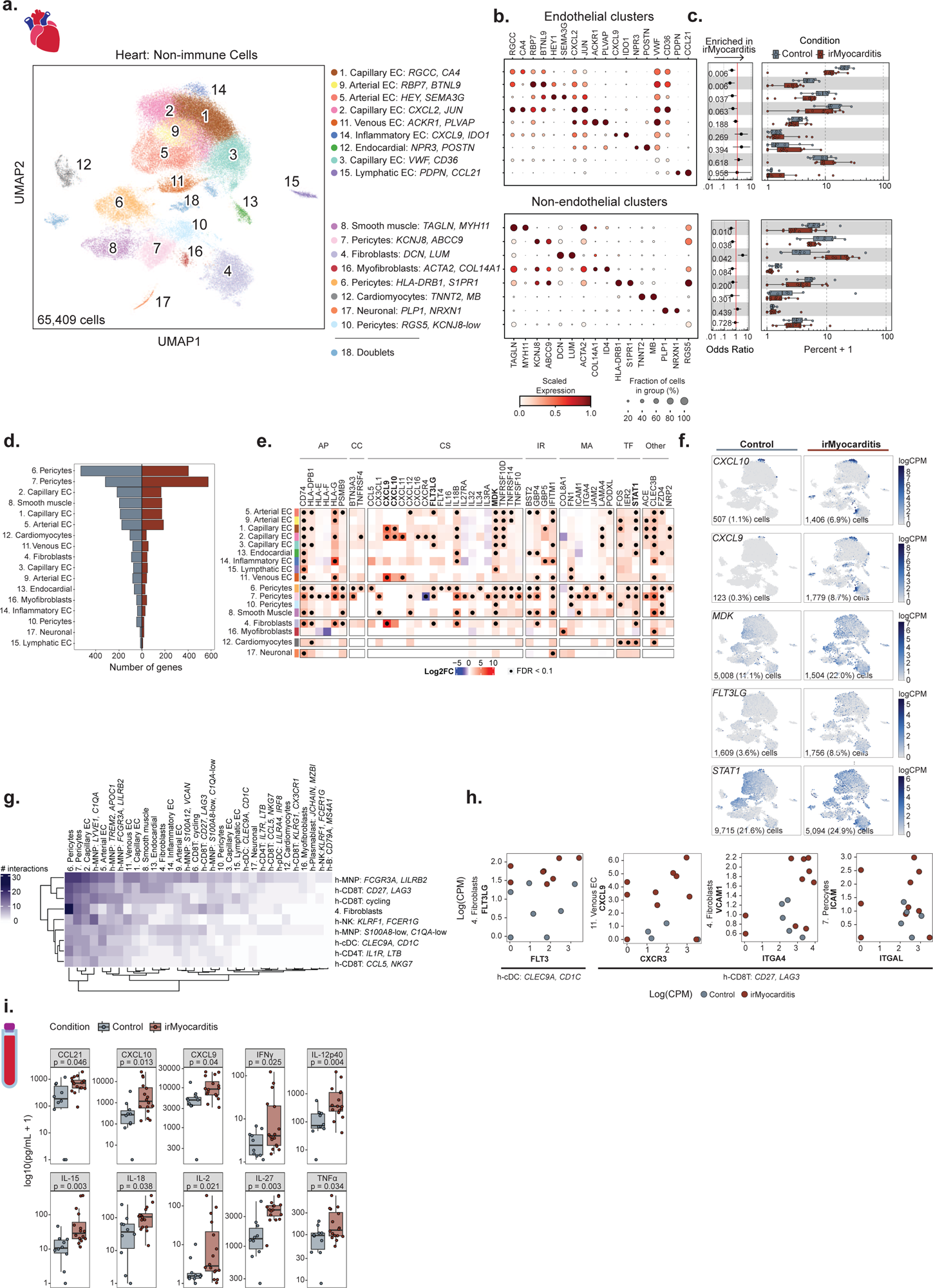
Immunomodulatory signals from non-immune intracardiac populations and circulating secreted factors nominate putative therapeutic targets for irMyocarditis. **a,** UMAP embedding of 65,409 non-immune cells isolated from the heart, segregated by endothelial cells (top) and non-endothelial cells (bottom), and colored by the 18 defined cell subsets labeled on the right. Cell subset number was assigned according to the absolute number of cells detected per subset and are listed in order of ascending p-value for the associated analyses in panel **c**. **b,** Dot plot showing top marker genes for each non-immune subset. Dot size represents the percent of cells in the subset with non-zero expression of a given gene. Color indicates scaled expression across subsets. **c,** Cell subset differential abundance analysis comparing irMyocarditis cases (n = 12, red) to control (n = 8, blue). Left: logistic regression odds-ratio for association of non-immune cell subset abundance with irMyocarditis (OR > 1) versus control (OR < 1) for each subset. Unadjusted likelihood-ratio test p-values for each subset are shown. Right: boxplots show per patient intracardiac non-immune cell subset compositions, where each dot represents a patient. Composition is reported as the percent of cells from a patient in each subset. **d,** The number of differentially expressed genes (DEGs) for each cell subset when comparing irMyocarditis cases (right, red) to control (left, blue). **e,** Heatmap showing select DEGs (irMyocarditis versus control) grouped by the following biological themes: antigen presentation (AP), co-inhibition or co-stimulation (CC), cytokine signaling (CS), interferon response (IR), motility and adhesion (MA), transcription factors (TF), and “Other” genes of biological interest. Color scale indicates log2 fold-change difference between irMyocarditis cases and controls. Black dots indicate FDR < 0.1 (Wald test). **f,** Feature plots using color to indicate gene expression (logCPM) levels of the indicated genes expressed by control samples (left) or irMyocarditis samples (right) projected onto the non-immune cell UMAP embedding. Cell numbers and percentages represent gene expression across respective control or irMyocarditis non-immune cells. **g,** Heatmap reporting the number of significant interactions between intracardiac cell subsets as predicted by CellphoneDB. Significant CellphoneDB results were filtered for high-confidence interactions (see **Methods**). **h,** Scatter plots showing select predicted receptor:ligand pairs of interest from CellphoneDB analysis. Each point represents a donor (red = irMyocarditis, blue = control) and the x- and y-axes represent the logCPM of receptors and ligands, respectively, from the indicated cell subset. **i,** Serum concentrations of selected cytokines and chemokines in irMyocarditis cases and controls. In **c**, error bars represent 95% confidence intervals. In **c** and **i**, boxes represent the median (line) and interquartile range (IQR) with whiskers extending to the remainder of the distribution, no more than 1.5x IQR, with dots representing individual samples.

### Stromal cell populations upregulate the expression of transcriptional programs involved in immune cell activation and recruitment

Differential gene expression analysis comparing irMyocarditis to control tissue demonstrated upregulation of 654 unique genes with FDR<0.1 in at least one stromal cell subset (**Figure 5d, Supplementary Table 14**). Unexpectedly, two pericyte subsets, one characterized by *HLA-DRB1* and *S1PR1* expression (cluster 6) and Pericytes^KCNJ8,ABCC9^ had the highest number of upregulated genes (396 and 563 genes, respectively). Multiple genes involved in antigen presentation, including *CD74* and *HLA-DPB1*, were upregulated across multiple non-immune cell subsets in irMyocarditis cases compared to controls (**Figure 5e**). The expression levels of numerous cytokines and chemokines were also increased in stromal cells across cases. For example, expression of some or all of the CXCR3 ligands *CXCL9, CXCL10,* and *CXCL11* were increased in each of the Capillary EC^RGCC,CA4^ (cluster 1), Capillary EC^CXCL2,JUN^ (cluster 2), venous EC, and fibroblasts subsets (**Figure 5e-f**). Midkine (*MDK*), a cytokine with a defined role in T cell activation^68^, was broadly upregulated across many endothelial and mural cell subsets. *FLT3LG,* the ligand for the FLT3 cytokine receptor and a key regulator of DC mobilization and function^51^, was increased across several EC subsets, mural cell subsets, and in fibroblasts. Expression of transcription factor *STAT1*, which is key in inducing the transcription of ISGs, was increased in multiple populations, analogous to observations in lymphoid and MNP populations.

### Cell-cell communication analysis nominates interactions involved in immune cell recruitment, activation, and retention to heart tissue

The interactions between immune and stromal cells in irMyocarditis represent a potentially under-explored opportunity to understand disease biology and identify therapeutic targets for mitigating irMyocarditis. We leveraged CellPhoneDB^69^ to explore intracellular interactions and focused on genes in ligand-receptor pairs where (a) one gene in the pair was differentially expressed in irMyocarditis heart tissue and (b) one cell subset expressing either the ligand or receptor was more abundant in irMyocarditis heart tissue (**Methods**). Results were manually curated for interactions that were supported by published literature to generate a final list of 92 unique receptor:ligand pairs and 1,069 subset-specific interactions (**Figure 5g**, **Supplementary Table 20**). Pericytes^HLA-DRB1,S1PR1^ and Pericytes^KCNJ8,ABCC9^ had the most predicted intercellular interactions, with a total of 116 interactions and 93 interactions across the eight cell populations enriched in irMyocarditis heart tissue, respectively. Endothelial subsets, fibroblasts, and MNP subsets each upregulated the expression of genes involved in the recruitment, activation, and retention of immune cells into irMyocarditis tissue (**Supplementary Table 14**). For example, *FLT3* expressed on h-cDCs^CLEC9A,CD1C^ is predicted to interact with *FLT3LG* expressed by fibroblasts and may be involved in the increased number of cDCs in irMyocarditis tissue, particularly in areas of active inflammation (**Figure 4c, 4f, 5f, 5h** far left). Stromal cell subsets may also play a role in the attracting and retaining T cells in heart tissue. For example, h-CD8T^CD27,LAG3^ cells are enriched in irMyocarditis and express the receptor *CXCR3* (**Figure 2c-d, 5h** left), which is also found on the circulating b-CD8T^cycling^ cells associated with fatal irMyocarditis **(Figure 3h-i**). Our analysis predicts that these CXCR3-expressing CD8T cells may interact with venous ECs and other stromal cell subsets expressing *CXCL9* (**Figure 5f**). h-CD8T^CD27,LAG3^ cells also express the integrins *ITGA4* (**Figure 2g, 5h** right) and *ITGAL* (**Figure 5h** far right), which are expected to interact with *VCAM1* on fibroblasts and *ICAM1* on Pericytes^KCNJ8,ABCC9^, respectively. These results suggest that the stromal cells in the heart may play an under-explored role in the pathophysiology of irMyocarditis.

### Increased levels of immunomodulatory factors are found in the blood of in irMyocarditis patients

Based on our results showing increased expression of genes encoding soluble inflammatory mediators in our MNP and stromal subsets in irMyocarditis, we investigated circulating levels of these immunomodulatory proteins. We analyzed serum from patients with irMyocarditis prior to treatment with corticosteroids (n = 16) and from ICI-treated patients without irAEs (n = 10) using a 71-analyte panel (**Methods**). Among these 71 examined factors, 16 were increased in the irMyocarditis samples (p < 0.05) (**Figure 5i, Supplementary Table 21**). These included CCL21 (also known as “6Ckine,” p = 0.046), which is known to recruit T cells, B cells, and DCs into lymphoid organs^70,71^. Chemokines CXCL9 (p = 0.040) and CXCL10 (IP10, p = 0.013), which mediate chemotaxis through their receptor CXCR3, were increased in serum of patients with irMyocarditis, and their transcripts were found to be upregulated in several non-immune cell populations (**Figure 5e-f**). Noteworthy, the receptor for these molecules, *CXCR3*, is expressed on b-CD8T^cycling^ cells (within which sharing of TCR clones found in irMyocarditis heart tissue correlated with fatal irMyocarditis) (**Figure 3f**), h-CD8T^CD27,LAG3^ cells, and h-CD8T^cycling^ cells (**Figure 2d**). Levels of TNFα (p = 0.034), which was also differentially expressed by h-MNP^TREM2,APOC1^ cells in irMyocarditis tissue (**Figure 4h**), IFN-γ (p = 0.025), and IL-12p40 (p = 0.004) were all upregulated in cases. Interestingly, these three cytokines are each targeted by agents that are commercially available and FDA-approved for the treatment of auto-immune diseases^72^. Additional factors with increased serum levels in irMyocarditis included IL-2 (p = 0.021), IL-15 (p = 0.003), IL-18 (p = 0.038), and IL-27 (p = 0.003), all of which support and direct T cell activation (**Figure 5i)**.

## Discussion

ICIs are a widely used class of cancer therapeutics that have demonstrated life-saving activity across a range of tumor types^1^, but they have the potential to cause morbid and potentially fatal irAEs, such as irMyocarditis^5–8^. Partly because severe irAEs may not be faithfully recapitulated in the murine tumor models used to study ICI efficacy, the pathogenesis of irMyocarditis and its relationship to anti-tumor immunity remains poorly characterized. We present the first systems-based approach and in-depth analysis of paired heart, blood, and tumor to report the first comprehensive, high-resolution study of the immune responses across tissue compartments that define irMyocarditis in patients. Our key findings are summarized in **Figure 6**. Our results indicate that irMyocarditis hearts are enriched for multiple CD8 T cell and MNP subsets, CD4 T cells, cDCs, and fibroblasts. Multiple transcriptional programs are activated in the setting of irMyocarditis, including those involved in interferon signaling, cell trafficking, antigen presentation, and immune cell activation. Sequencing of TCRs demonstrated that the clones most enriched relative to control tissue in the heart and in autologous tumors were largely non-overlapping, suggesting that distinct antigens drive T-cell activity in these two tissues. Multiple novel biomarkers of irMyocarditis severity were identified: heart-expanded T cell clones disproportionately shared with circulating b-CD8T^cycling^ cells in fatal cases and correlations between intracardiac h-cDCs^CLEC9A,CD1C^ and h-CD8T^cycling^ populations and serum troponin. Multiple soluble factors were found at increased levels in the serum of irMyocarditis patients, including several targeted by approved therapeutics.

**Figure 6.**
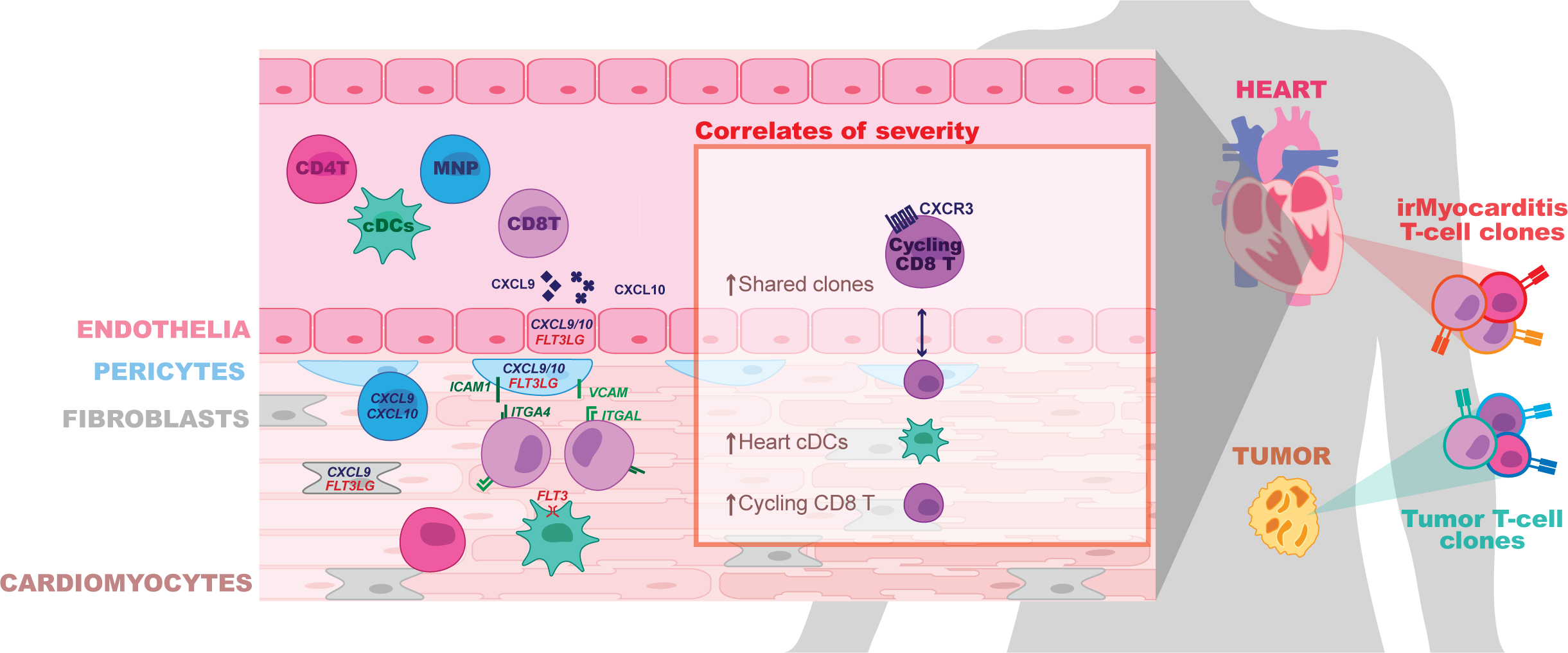
Summary of key findings. The key findings of our study are represented, starting on the left, by showing the increased cardiac infiltration of CD4 T cells (CD4T), CD8 T cells (CD8T), mononuclear phagocytes (MNP), and conventional dendritic cells (cDCs). These immune cells are shown expressing some of the genes (*ITGA4* and *ITGAL* on CD8T cells; *CXCR3* on cycling CD8T; *FLT3* on cDCs) that are anticipated to be involved in the recruitment and retention of immune cells into the heart by cognate ligands expressed by other cells in the heart (*CXCL9/10* and *FLT3LG* in endothelial cells; CXCL9/10, *ICAM1*, and *VCAM* in pericytes; *CXCL9/10* in MNPs; *CXCL9* and *FLT3LG* on fibroblasts). Secreted CXCL9 and CXCL10 are highlighted as proteins that are elevated in the blood of irMyocarditis patients. “Correlates of severity” highlight factors in our scRNA-seq analysis that are associated with fatality (increased sharing of TCR clones expanded in the heart with cycling CD8T cells in the blood) and increased serum troponin in irMyocarditis patients (increased cDCs and cycling CD8T in the heart). An illustration of the heart and paired tumor highlight T cells with distinct colors to emphasize the finding that different T-cell clones are enriched in irMyocarditis and tumor compared to controls.

While we were able to discern clear and consistent biological signals across all patients in our study, the patchy nature of the disease, tissue availability, and limited ICI-treated controls are limitations of our study. irMyocarditis is histologically heterogeneous, often with relatively small regions of inflammation interspersed within otherwise normal-appearing myocardium^12,19,20,73^. The distinction between irMyocarditis cases and controls in our scRNA-seq analysis was based on the histologic finding of irMyocarditis in that patient, confirmed by a pathologist; however, endomyocardial biopsies provide scant tissue that needed to be processed immediately and in its entirety to generate scRNA-seq data, which makes it impossible to determine how the tissue used for scRNA-seq captured the patchy nature of irMyocarditis. Autopsy samples provide ample tissue, but these patients are almost always heavily treated with immunosuppressive agents that alter gene expression profiles. Additionally, since biopsies are typically performed only in the setting of high clinical suspicion and with evidence of cardiac dysfunction, ICI-treated control samples are rarely available, have few immune cells, and may have non-inflammatory cardiac disease. We leveraged a published heart atlas^29^ to account for some of these factors, but we were underpowered to compare fatal to non-fatal irMyocarditis tissue samples and may not be able to detect various endotypes of irMyocarditis.

Despite these limitations, our study recapitulates findings from prior histopathological analyses^19,20^ and builds a more detailed cellular and molecular understanding of irMyocarditis in cancer patients. Prior work has shown that irMyocarditis tissue is characterized by infiltrating T cells and macrophages^19,20^. Beyond validating the enrichment of these populations, our scRNA-seq analysis further defined the cellular phenotypes of the enriched CD4 T cells, cytotoxic CD8 T cells, and multiple MNP subsets in irMyocarditis heart tissue. Additionally, we identified an increase in cDCs and found that the presence of h-cDCs^CLEC9A,CD1C^ and h-CD8T^cycling^ correlate with circulating troponin, a clinical measure of myocardial damage. Our study is also the first to deeply profile stromal cells in irMyocarditis. We found that endothelial, fibroblast, and pericyte populations have increased expression of genes that could play a role in the recruitment, activation, and retention of immune cells into the heart. Collectively, our data supports a pathophysiologic model of irMyocarditis in which potentially auto-reactive cytotoxic CD8 T cells and antigen presenting cells are both recruited to and retained in the heart by stromal and immune-cell signaling networks.

The identity of the antigen (or antigens) driving cardiac T-cells responses in irMyocarditis remains unclear, as is whether the same antigen drives anti-tumor immunity in these patients. A seminal case report of irMyocarditis found common expanded TCR sequences in irMyocarditis tissue and tumor^19^, supporting the potential for shared antigens in tumor and irAEs that has been subsequently explored in irAEs affecting the skin^74,75^ and lungs^76^. It is possible that irMyocarditis may be pathophysiologically or antigenically heterogeneous; indeed, some of our irMyocarditis cases had evidence of clinically occult cardiac metastases in biopsy or autopsy samples, which could play a role in initiating, sustaining, or mimicking irMyocarditis. However, in each of our four cases without diffuse cardiac metastases, we utilized control tissues to account for potentially ubiquitous, non-specific “bystander” TCR clones^45^, and we found that the TCR clonotypes most enriched in hearts and autologous tumors relative to control tissues were distinct. Though the antigens recognized by these TCRs are presently unknown and disparate TCR sequences can be specific for the same antigen, our findings of different enriched clonotypes in the heart and tumor within each patient argue against a common antigen driving the immune responses at both sites. The cardiac protein α-myosin (encoded by *MYH6*) has been reported as an auto-antigen in mouse models of irMyocarditis^23,25^, and α-myosin-specific T cells are also found in the blood and heart of irMyocarditis patients^23^. Notably, however, α-myosin-reactive T-cell clones were reported in the blood of every healthy control donor and were not the most expanded clones recovered from irMyocarditis tissue. Further investigation into antigenic drivers and mediators of inflammation in irMyocarditis and tumor are needed to better define the relationship between T-cell responses in these different microenvironments. These studies may also clarify why only approximately 1% of ICI recipients develop irMyocarditis despite the high prevalence of T-cell clones specific for α-myosin^23^, which could help to identify cancer patients at risk for developing irMyocarditis.

Across our cohort, the analysis of blood and paired tissue yielded insights into the pathology of irMyocarditis and nominated biomarkers that could be used to design new diagnostic and therapeutic strategies. We observed a positive correlation between MNP cells in blood serum troponin with a corresponding inverse correlation between serum troponin and the frequencies of CD4 T cells, CD8 T/NK cells, and B/plasma cells. Patients with fatal irMyocarditis were also more likely to share heart-expanded TCR clonotypes within a circulating, CD8 T population expressing *CXCR3* and *MKI67* (b-CD8T^cycling^). Secreted factor analysis found that irMyocarditis patients had increased levels of chemokines (CCL21, CXCL9, and CXCL10), targetable inflammatory mediators (TNFα, IFN-γ, and IL-12p40), and T-cell supporting cytokines (IL-2, IL-15, IL-18, and IL-27) compared to ICI-treated controls.

Our efforts also help to contextualize irMyocarditis within the emerging literature of irAEs impacting other organs and distinct inflammatory conditions impacting the heart. While irAEs in barrier organs, such as the colon, may be driven by both resident and recruited immune cells^35,77^, it appears as though irAEs in sterile organs such as the heart and joints are driven by immune cell recruitment^78^. Many efforts to study irAEs across affected tissues have implicated CXCR3 and its ligands CXCL9/10/11 in the recruitment of immune cells into various tissues^21,35,77–80^. This chemokine system is important for anti-tumor responses in the tumor microenvironment^81^, and while these ligands are increased in the blood of all patients receiving ICIs^82^, it is tempting to speculate that the degree of elevation can distinguish irAEs and even the severity of irAEs, such as fatal irMyocarditis. Despite these similarities, differences are observed in organ-specific irAE studies, such as Th17 signals reported in irArthritis and irColitis that are absent in our data^35,78^. Organ-specific differences may also explain why some immunosuppressive agents may be helpful in some irAEs but harmful in others; for instance, TNF-α inhibitors have shown efficacy in irColitis but may be detrimental in irMyocarditis^83^. We are the first to show the direct presence of cDCs in irAEs, but RNA expression profiles and secreted factors observed in ICI-related nephritis suggest that DCs and tertiary lymphoid structures may play a role in other irAEs as well^80^.

Despite the insights offered by this study, many gaps in knowledge remain regarding the underlying biology of irMyocarditis. Additional mechanistic, translational, and clinical research will be needed to identify optimal strategies to diagnose, risk-stratify, and manage patients with irMyocarditis. Validation of the candidate biomarkers for irMyocarditis severity suggested herein may be useful towards this goal. The ideal treatment for irMyocarditis would minimize the cardiac damage while also allowing anti-tumor benefits of ICIs to persist. Future studies of agents used to treat irMyocarditis should therefore assess if they also blunt^16,17,84,85^, permit^86–88^, or even stimulate anti-tumor responses^89^. This framework could also be applied broadly to irAEs impacting other organ systems. Identifying patients at risk and disentangling irAEs from effective anti-tumor immune responses will ultimately help to maximize the benefit of ICIs while minimizing the harm from irAEs.

## Methods

### Study design, patient identification, and sample selection – heart specimens for scRNA-seq

Heart tissue was collected at our institution through endomyocardial biopsy or autopsy of patients receiving ICI agents for the treatment of cancer. Informed consent was obtained from all patients or their appropriate representatives. All research protocols were approved by the Dana-Farber/Harvard Cancer Center Institutional Review Boards (#11-181 and 13-416). Endomyocardial biopsies were collected in the catheterization laboratory as part of the clinical evaluation for suspected irMyocarditis, and a single tissue fragment (∼1-2 mm^3^) primarily derived from the right side of the interventricular septum was collected for research. One patient (SIC_264) had a biopsy of the left ventricle for clinical reasons. Autopsy samples were obtained by an autopsy pathologist from the right ventricular free wall. Autopsy samples were obtained rapidly (within 6 hours) following the death of patients on ICI agents with or without clinically suspected irMyocarditis. Patients were classified as irMyocarditis cases based on pathological review of tissues by specialized cardiac pathologists as part of routine clinical care. Control heart samples were obtained through two sources: (i) Two samples (SIC_182 and SIC_333) were obtained at our institution from cancer patients who received ICIs and underwent endomyocardial biopsies and/or autopsy with a clinical suspicion for irMyocarditis, but they ultimately did not have histopathological features of irMyocarditis. (ii) Six additional controls were obtained from hearts that were offered but not selected for transplantation and were used to generate a scRNA-seq heart atlas, as described^29^; none of these patients were known to be on ICI therapy. To account for differences in cell proportions or gene expression that occurs in different regions of the heart^90^, we used samples from this control cohort that were obtained from the right ventricle septum or right ventricular free wall to match the anatomic location of samples collected at our institution. The heart atlas also included samples that were enriched for CD45^+^ cells, which we included in our clustering and select downstream analyses, as outlined in **Figure 1b**.

### Study design, patient identification, and sample selection – heart and tumor specimens for bulk TCR-β sequencing

Samples for bulk T-cell receptor β chain (TCR-β) sequencing were identified from the autopsies of cases enrolled in our scRNA-seq cohort as well as additional patients who consented to our tissue banking protocols and were found to have histological evidence of irMyocarditis at time of autopsy. Control heart tissues for bulk TCR-β sequencing were identified from patients who were receiving an ICI, did not have evidence of irMyocarditis at autopsy, and for whom archival tissue was available through our biobanking protocol. Tissues were annotated by a cardiac pathologist as “active,” “borderline,” or “healing” irMyocarditis, in accordance with the Dallas Criteria^43^.

### Study design, patient identification, and sample selection – blood and serum specimens

Paired blood and serum were sought prior to the initiation of corticosteroids for the treatment of irMyocarditis in our tissue scRNA-seq cohort as well as additional patients who were diagnosed with irMyocarditis but did not have tissue data. Where available, blood samples were collected or accessed from collaborating biobanking efforts at the following clinically relevant timepoints: prior to the start of ICI (“pre ICI” timepoints); after the start of ICI but without clinical irMyocarditis (“on ICI”); at the time of clinical irMyocarditis diagnosis but prior to steroid initiation (“pre-corticosteroid”); after steroid initiation or subsequent therapies (collectively, “post-corticosteroid” samples). Control serum samples were identified from biobanked specimens from patients receiving an ICI who did not have evidence of irMyocarditis or any other active irAE.

### Clinical covariates

Clinical data were obtained retrospectively from electronic medical records, including patient demographics, cancer type, prior cancer therapies received, specific ICI agents used, method of irMyocarditis diagnosis, myocardial biopsy pathology grade, troponin T measurements, and concomitant irAEs.

### Preparation of tissue samples for scRNA-seq

Tissue samples obtained by biopsy or autopsy were immediately placed in ice-cold HypoThermosol solution (BioLife Solutions, Bothell, WA) and kept on ice during transfer to the research facility. Tissue was then washed twice with cold phosphate buffered saline prior to being dissociated with the human tumor dissociation kit according to the manufacturer’s instructions (Miltenyi Biotic, Bergisch Gladbach, Germany), with modification such that calcium chloride was added to the enzymatic cocktail to a final concentration of 1.25 mM. Biopsies were cut into ≤ 1 mm pieces with standard laboratory tissue dissection scissors. Tubes containing tissue fragments in the enzymatic cocktail were placed in a heated shaker at 37° C with shaking at 750 RPM for 25 minutes with the machine placed on its side to prevent tissue fragments from settling. Following incubation, the reaction was quenched through the addition of 100 µl human serum. The mixture was further dissociated through manual trituration followed by filtration through 70 µm mesh. Following centrifugation at 350 x g for 12 minutes, the supernatant was removed, and RBC lysis was performed for two minutes (ACK lysing buffer, Lonza, Basel, Switzerland). Following a wash step, cells were resuspended in phenol-free RPMI with 2% (v/v) human AB serum.

Due to excessive debris, the following myocardial samples obtained from autopsy were sorted by FACS for live, non-red blood cells following tissue dissociation; SIC 176, SIC 182_B, SIC 232, and SIC 264_B (B samples refer to the second of two samples when collected from the same patient). Tissue was prepared for cell sorting by first undergoing dissociation as described above. Following the resuspension of cells in sort buffer (phenol-free RPMI with 2% [v/v] human AB serum), Fc receptors were blocked (Human TruStain FcX, Biolegend 422302), after which cells were incubated with CD235a-PE-Cy5 (Biolegend 306606) for 30 minutes. Following a wash, cells were resuspended in sort buffer containing DAPI. Flow cytometric sorting was performed on a Sony MA900 Cell Sorter (Cell Sorter Software v 3.3.0) to collect live, singlet, CD235a^-^ cells (**Supplementary Figure 9**). Sorted cells were centrifuged and resuspended in sort buffer prior to loading in the 10x Chromium chip.

### PBMC CD45^+^ enrichment, cell hashing, and CITE-seq staining

Cryopreserved PMBC samples (total 57 samples across multiple timepoints from 27 patients; **Supplementary Tables 4-6**) were thawed at 37°C, diluted with a 10x volume of RPMI with 10% heat-inactivated human AB serum (Sigma), and centrifuged at 300 x g for 7 minutes. Cells were resuspended in CITE-seq buffer (RPMI with 2.5% [v/v] human AB serum and 2mM EDTA) and added to 96-well plates. Dead cells were removed with an Annexin-V-conjugated bead kit (Stemcell 17899) and red blood cells were removed with a glycophorin A-based antibody kit (Stemcell 01738); modifications were made to manufacturer’s protocols for each to accommodate a sample volume of 150 uL. Cells were quantified with an automated cell counter (Bio-Rad TC20), after which 2.5 x 10^5^ cells were resuspended in CITE-seq buffer containing TruStain FcX blocker (Biolegend 422302) and MojoSort CD45 Nanobeads (Biolegend 480030). Hashtag antibodies (Biolegend) were added to samples (**Supplementary Table 11**) followed by a 30 minute incubation on ice. Cells were then washed three times with CITE-seq buffer using a magnet to retain CD45^+^ cells. Live cells were counted with trypan blue, and 7-8 samples (each sample with 60,000 cells) bearing different hashtag antibodies were pooled together at equal concentrations. Pooled samples were filtered with 40 μM strainers, centrifuged, and resuspended in CITE-seq buffer with TotalSeq-C antibody cocktail (Biolegend; **Supplementary Table 11**). Cells were incubated on ice for 30 minutes, followed by three washes with CITE-seq buffer and a final wash in the same buffer without EDTA (RPMI with 2.5% [v/v] human AB serum). Cells were resuspended in this buffer without EDTA, filtered a second time, and counted.

### scRNA-seq data generation

For heart samples, single cell suspensions containing up to 12,000 cells (or all available cells when total was <12,000) were loaded per channel in 10x Chromium chips. For some samples, two wells in the 10x Chromium chip were loaded to maximize cell recovery; downstream data from these multiple wells were later combined and considered as a single sample. For hashed PBMC samples, samples were diluted to a concentration of 1,200 cells/μl, and 50,000 cells were loaded per well into the 10x Chromium chip channel. According to the manufacturer’s instructions, cell/bead emulsions were generated and transferred to PCR tube strips, after which cDNA libraries were generated. Libraries from heart samples were generated using Chromium Single Cell 5’ V1 kits (10x Genomics PN-1000006) except for one sample (SIC_317), which was generated with a 5’ V2 kit (10x Genomics PN-1000263) (**Supplementary Table 2)**. Hashed PBMC single cell libraries were generated with the Chromium Single Cell 5’ kit (V1.1, 10x Genomics PN-1000020) together with the 5’ Feature Barcode library kit (10x Genomics PN-1000080) (**Supplementary Table 5**). TCR-enriched cDNA libraries were generated with the Chromium Single Cell V(D)J Enrichment kit (10x Genomics PN-1000005). Library quality was assessed with an Agilent 2100 Bioanalyzer.

All heart sample gene expression libraries were sequenced on an Illumina Nextseq 500/550 instrument using the high output v2.5 75 cycles kit with the following sequencing parameters: read 1 = 26; read 2 = 56; index 1 = 8; index 2 =0. TCR-enriched libraries from heart tissue were sequenced on an Illumina Nextseq 500/550 instrument using the high output v2.5 150 cycle kit with the following sequencing parameters: read 1= 30, read 2 = 130, index 1 = 8, index 2 = 0. All hashed PBMC CITE-seq, feature barcode (i.e., ADT and HTO libraries), and TCR libraries were sequenced on an Illumina Novaseq instrument using the S4 300 cycles flow with the following sequencing parameters: read 1 = 26; read 2 = 91; index 1 = 8; index 2 =0.

### scRNA-seq read alignment and quantification

Raw sequencing data was pre-processed with CellRanger (v3.0.2, 10x Genomics) to demultiplex FASTQ reads, align reads to the human reference genome (GRCh38, v3.0.0 from 10x Genomics), and count unique molecular identifiers (UMI) to produce a cell x gene count matrix^91^. All count matrices were then aggregated with Pegasus (v1.1.0, Python) using the aggregate_matrices function^92^. Low-quality droplets were filtered out of the matrix prior to proceeding with downstream analyses using the percent of mitochondrial UMIs and number of unique genes detected as filters (heart tissue = < 20% mitochondrial UMIs, > 300 unique genes; PBMCs = < 20% mitochondrial UMIs, > 400 unique genes). The percent of mitochondrial UMI was computed using 13 mitochondrial genes (MT-ND6, MT-CO2, MT-CYB, MT-ND2, MT-ND5, MT-CO1, MT-ND3, MT-ND4, MT-ND1, MT-ATP6, MT-CO3, MT-ND4L, MT-ATP8) using the qc_metrics function in Pegasus. The counts for each remaining cell in the matrix were then log-normalized by computing the log1p(counts per 100,000), which we refer to in the text and figures as logCPM. The detailed quality control statistics for these datasets are compiled in **Supplementary Tables 2 and 6**.

### Basic clustering

All cells from all samples were utilized for clustering. First, 2,000 highly variable genes were selected using the *highly_variable_features* function in Pegasus and used as input for principal component analysis. To account for technical variability between donors, the resulting principal component scores were aligned using the Harmony algorithm^93^. The resulting principal components were used as input for Leiden clustering and Uniform Manifold Approximation and Projection (UMAP) algorithm (spread=1, min-dist=0.5). The number of principal components used for each blood clustering (CD4 T = 16; CD8 T/NK = 18; MNP = 24) and tissue clustering (T/NK = 24; MNP = 35; non-immune = 50) was decided via molecular cross-validation^94^.

### Marker gene identification and cell annotation

The marker genes defining each distinct cell subset from our global and lineage-specific subclustering analyses were determined by applying two complementary methods. First, we calculated the area under the receiver operating characteristic (AUROC) curve for the logCPM values of each gene as a predictor of cluster membership using the *de_analysis* function in Pegasus. Genes with an AUROC ≥ 0.75 were considered marker genes for a particular cell subset. Second, we created a pseudobulk count matrix^95^ by summing the UMI counts across cells for each unique cluster/sample combination, creating a matrix of n genes x (n samples*n clusters). We performed “one-versus-all” (OVA) differential expression (DE) analyses for each cell subset using the Limma package (v3.54.0, R)^96^. For each subset, we used an input model *gene ∼ in_clust*, where *in_clust* is a factor with two levels indicating if the sample was in or not in the subset being tested. A moderated T-test was used to calculate P values and compute a false discovery rate (FDR) using the Benjamini-Hochberg method. We identified marker genes that were significantly associated with a particular subset as having an FDR < 0.05 and a log2 fold change > 0. The AUROC and OVA pseudobulk marker genes for all cell subsets can be found in **Supplementary Tables 8 and 12**. Marker genes for each cell subset were interrogated and investigated in the context of other published immune profiles to guide our cell subset annotations.

### Abundance analysis

To identify the association between cell subset abundance and patient group (irMyocarditis, control), we used a mixed-effects association logistic regression model similar to that described by Fonseka et al.^97^. We used the *glmer* function from the lme4 package (v1.1-31, R) to fit a logistic regression model for each cell subset. Each subset was modeled independently with a “full” model as follows:

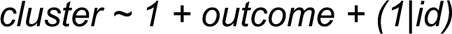

where *cluster* is a binary indicator set to 1 when a cell belongs to the given cell subset or 0 otherwise, *outcome* is a factor with 2 levels (irMyocarditis or control), and *id* is a factor indicating the donor. The notation *(1|id)* indicates that *id* is a random intercept. To determine significant associations, a “null” model of *cluster ∼ 1 + (1|id)* was fit and a likelihood ratio test was used to compare the full and null models. A false discovery rate was calculated using the Benjamini-Hochberg approach and clusters with an FDR < 10% were considered significant.

Since we were underpowered to investigate differences between fatal and non-fatal irMyocarditis in our heart scRNA-seq data, we sought to identify associations between cell subset frequencies and clinically measured serum troponin levels, a previously described correlate of irMyocarditis severity^5,36^. We performed linear regression on each cell subset individually, using the model *log(troponin) ∼ log(abundance)* where *troponin* is an individual’s serum troponin T level measured closest to time of tissue sample collection (no greater than ±2 days) and *abundance* represented the relative proportion a cell subset represented in a given sample.

To accurately measure the abundance of each cellular component of heart tissue, we used only unenriched samples (“native” fractions) and excluded the CD45^+^-enriched samples from the published heart atlas^29^. In the case of SIC_182, a control patient who underwent a biopsy (sample “SIC_182_A”) and then an autopsy (“SIC_182_B”) but did not have irMyocarditis and never received immunosuppression, data from this patient’s two samples were combined and treated as a single timepoint.

### Differential gene expression analysis

Comparisons between irMyocarditis and control heart tissue was limited to just samples prior to corticosteroids administration (“pre-corticosteroid”) and included both native and CD45^+^ enriched samples from the heart cell atlas to account for the paucity of immune cells in uninflamed hearts^29^ (**Figure 1B**). Analyses were performed on pseudobulk count matrices using the DESeq2 package (v. 1.38.2, R). The input model was *gene ∼ case,* where *case* was a binary variable indicating if the sample came from an irMyocarditis patient or control.

Significant differentially expressed genes (DEGs) were identified using a Wald test (FDR < 0.1) (**Supplementary Table 14**). Select DEGs were visualized with the ComplexHeatmap package (v.2.14.0, R).

### Cell-cell communication analysis

To infer potential receptor:ligand interactions between cell-cell pairs, we used CellPhoneDB (v2.1.7, Python) and ran the algorithm independently on irMyocarditis heart and control cells. Each cell subset was tested as both a sender (ligand) and receiver (receptor) population as defined by the algorithm, and all possible combinations of cell-cell pairs were tested. We restricted potential interactions to those where the receptor and ligand were each expressed in >10% of their respective cell subset and in at least 20 cells, with significance defined as an empirical P < 0.001. These potential interactions were further filtered to include only those where at least the receptor and/or ligand were differentially expressed between irMyocarditis and control (see above *Differential Gene Expression* section), and the receptor:ligand interaction was validated by an exhaustive literature search.

### Secreted factors analysis

Peripheral blood was collected from irMyocarditis and control patients in serum separator tubes. Following centrifugation for 10 minutes at 890g, serum was collected and stored at -80° C. Aliquots of undiluted serum were analyzed for the presence of 71 secreted factors through multiplex immunoassay (Human Cytokine Array/Chemokine Array 71-Plex Panel [catalog #HD71], Eve Technologies, Calgary, AB, Canada). Log +1 transformed values were compared using a two-sided T-test.

### Tissue/bulk TCR data generation

Four autopsy cases were identified with available matched irMyocarditis, tumor, and histologically normal tissue adjacent to tumor. This normal adjacent tissue was used as “control” tissue. Formalin-fixed, paraffin-embedded (FFPE) slides were stained for hematoxylin and eosin (H&E) and annotated by a board-certified cardiac pathologist to indicate regions of irMyocarditis, tumor, and normal parenchyma. Marked regions of interest were manually macroscopically dissected using a scalpel to scrape tissue from serial unstained slides into 1.5 ml Eppendorf tubes pre-filled with 1 ml of xylene. Genomic DNA was extracted using the AllPrep DNA/RNA FFPE Kit instructions (Qiagen). T-cell receptor β chain (TCR-β) sequencing of the CDR3 region was performed (immunoSEQ ® human T-cell receptor beta [hsTCRB], Adaptive Biotechnologies)^42^. We also profiled heart tissue for four cancer patients receiving an ICI who consented to our collection protocol, underwent an autopsy, and did not have evidence of irMyocarditis (“heart controls”) (**Supplementary Table 3**).

### Comparison of irMyocarditis and tumor TCR-β repertoires

For comparisons between heart and tumor TCR repertoires, the frequencies, and not the absolute counts, of TCR clones were compared to account for differential recovery of TCR-β sequences from various tissues^98^. Expanded TCR-β sequences were defined as those accounting for >0.5% of all TCR-β sequences recovered from the tissue of interest (heart or tumor). These expanded sequences were then filtered to exclude bystander TCR-β sequences by performing a Fisher’s exact test on each, comparing the proportion of the TCR-β sequence in adjacent normal tissue to that of the tissue of interest (heart or tumor). Sequences with an FDR < 5% were considered enriched in their given tissue site (heart or tumor).

### TCR-β sequence diversity

Diversity curves that measured Hill’s diversity metric across diversity orders 0–4 were created using the package *alakazam* (v1.0.2, R) with the *alphaDiversity* function^99^. Hill’s diversity metric was only calculated on samples with ≥100 total TCR-β sequences.

### Examination of shared TCR-β clones in heart and blood

For bulk TCR-β-seq data, expanded TCR-β sequences were defined as those that represented > 0.5% of the bulk repertoire and were found to have a read count ≥ 2. For sc-TCR-β-seq data, expanded TCR-β sequences were those that represented > 0.5% of the sc-TCR-βseq repertoire and were found in ≥ 2 cells. For patients with both bulk TCR-β-seq and sc-TCR-β-seq data, these data were analyzed together. Expanded heart TCR-β clones were then compared to TCR-β clones recovered from PBMC scRNA-seq data. PBMC cell subsets were investigated for their association with the presence of expanded TCR-β heart clones by fitting a logistic regression model using the *glmer* function from the lme4 package (v1.1-31, R). For each donor, each subset was modeled independently with the full model *cluster ∼ 1 + myo_clone,* where *myo_clone* is a binary variable where the value was 1 if the cell had an expanded irMyocarditis TCR-β and 0 if it did not. An alternative null model *cluster ∼ 1* was then fit and a likelihood ratio test was used to compare the full and null models. Cell subsets with an FDR < 5% were considered to be enriched for expanded irMyocarditis TCR-β sequences.

### CD1c Staining

FFPE tissue sections from six irMyocarditis cases and four control hearts were stained using a Leica Bond RX automated stainer. Anti-CD1c-OTI2F4 mAb (Abcam ab156708) were labeled with DAB chromogen (Leica Bond Polymer Refine Detection DS9800) using EDTA based pH 9 epitope retrieval condition and 20 ug/ml antibody concentration.

### Image processing/data analysis

Whole slide images of stained slides were acquired at 40x (0.13um/pixel) resolution using an MoticEasyScan Infinity digital pathology scanner. Cell segmentation and phenotyping was performed within annotated tissue regions using HALO Image Analysis Platform (Indica Labs, v3.32541.184). Areas that contained debris or tissue folds were excluded from the analysis. The Multiplex IHC v3.1.4 module was used for identification of putative CD1c^+^ cells, which was based on signal intensity within the nuclear and cytoplasmic compartments. The candidate cells suggested by this automated analysis were then manually validated for each sample for cell phenotyping quality. Summary tables containing cell phenotype information were exported for analysis across samples (**Supplementary Table 19**).

### Data and Material availability

scRNA-seq count matrices and related data will be available in the GEO database upon publication, and raw human sequencing data will be available in the controlled access repository dbGaP (https://www.ncbi.nlm.nih.gov/gap/) upon publication.

### Code availability

Source code for data analysis will be available on GitHub upon publication of this study. A full list of software packages and versions included in the analyses is included in **Supplementary Table 23**.

## Acknowledgement

We are deeply grateful to all donors and their families. We also thank the Mass General Cancer Center, Ellison 16, and the Severe Immunotherapy Complications Service for their collaboration and support. S.M.B was supported by a National Institutes of Health T32 Award (2T32CA071345-21A1; PI Haber) and SITC-Mallinckrodt Pharmaceuticals Adverse Events in Cancer Immunotherapy Clinical Fellowship. D.A.Z. was supported by a National Institutes of Health T32 Award T32HL007208 (PI: Rosenzweig) and K24HL150238-02 (PI: Neilan). L.Z. was supported by the Spanish Society of Medical Oncology (SEOM) grant for a 2-year translational project at the MGH Cancer Center. K.S. was supported by a NIAID grant T32AR007258. P.S. is supported by the National Institutes of Health K08 Award (NHLBI K08 HL157725) and American Heart Association Career Development Award. M.F.T. is supported by the National Institutes of Health K08 Award (NIDDK K08 DK127246). G.M.B. is supported by an Adelson Foundation award. T.G.N is supported by a gift from A. Curt Greer and Pamela Kohlberg and from Christina and Paul Kazilionis, the Michael and Kathryn Park Endowed Chair in Cardiology, a Hassenfeld Scholar Award, and has additional grant funding from the National Institutes of Health/National Heart, Lung, and Blood Institute (R01HL137562, K24HL150238, R01HL130539). This work was made possible by the generous support from the National Institute of Health Director’s New Innovator Award (DP2CA247831; to A.C.V.), the Massachusetts General Hospital Transformative Scholar in Medicine Award (to A.C.V.), the Damon Runyon-Rachleff Innovation Award (to A.C.V.), The Melanoma Research Alliance Young Investigator Award (https://doi.org/10.48050/pc.gr.143739; to A.C.V.), the MGH Howard M. Goodman Fellowship (to A.C.V.), the Arthur, Sandra, and Sarah Irving Fund for Gastrointestinal Immuno-Oncology (to A.C.V.), the Kraft Foundation Award (to. K.L.R. and A.C.V.), and by the generous support of an anonymous donor (to. K.L.R. and A.C.V.).

## Author Contributions

S.M.B, D.A.Z., M.F.T., K.L.R., T.G.N, and A.C.V. conceived of and led the study; S.M.B, D.A.Z., M.F.T., and A.C.V. led the experimental design; S.M.B, D.A.Z., M.F.T., and P.S. carried out experiments with assistance from A.T., K.M., J.T., B.Y.A., J.B, N.S, S.C.M, and J.M.; N.P.S, I.K., S.R., and M.N. designed and performed computational analysis; S.M.B, D.A.Z., Y.S., and L.T.N. designed and performed microscopy experiments; S.M.B, D.A.Z., L.Z., P.C., C.J.P., D.M., M.J., J.F.G., D.J., M. M-K, R.J.S., G.M.B., J.S., T.G.N., K.L.R. provided clinical expertise, coordinated and performed sample acquisition, and/or established research protocols; M. M-K. and J.S. provided histopathology expertise; J.C. performed endomyocardial biopsies; A.C.V. managed and supervised the study; A.C.V., T.G.N., K.L.R., S.M.B., and D.A.Z. provided funding for this work; S.M.B, D.A.Z., N.P.S, I.K., and A.C.V. wrote the manuscript, with input from all authors.

## Supplementary Figure Legends

**Supplementary Figure 1.**
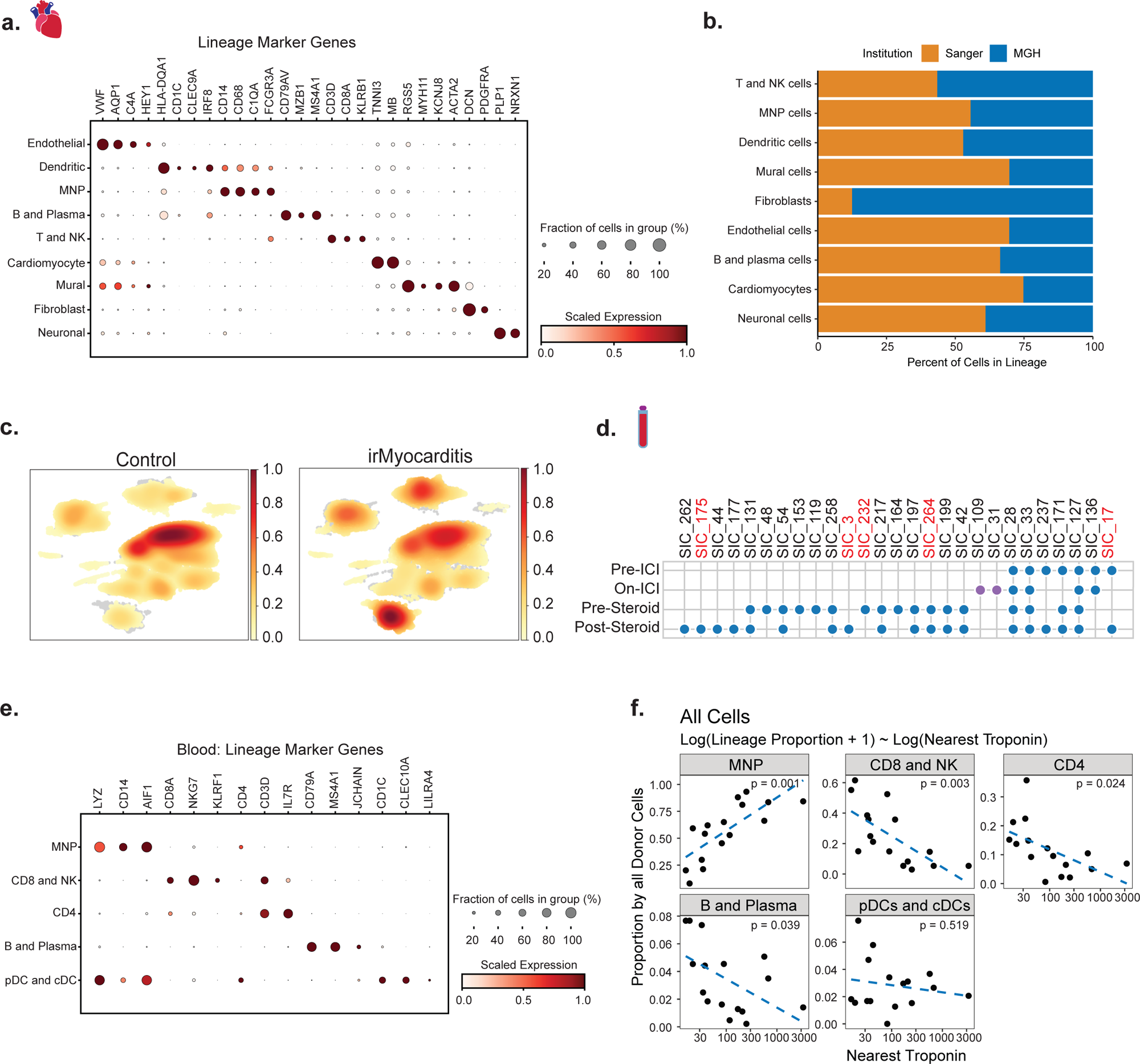
Cell lineages in heart and blood defined by scRNA-seq. **a,** Dot plot showing top marker genes for each lineage in the heart. Dot size represents the percent of cells in the lineage with non-zero expression of a given gene. Color indicates scaled expression across lineages. **b,** Stacked bar plots showing the composition of major cell lineages, colored by data source (“MGH” refers to data generated in this study; “Sanger’’ refers to public heart atlas data)^29^. **c,** UMAP embedding of cell density plot displaying the relative proportion of cells from irMyocarditis cases and controls. **d,** Blood samples and timepoints collected for the study. Red patient labels denote samples from patients with fatal irMyocarditis. Two ICI-treated patients who did not develop irMyocarditis (SIC_31, SIC_109) are denoted with purple dots. **e,** Dot plot showing top marker genes for each lineage in the blood. Dot size represents the percent of cells in the lineage with non-zero expression of a given gene. Color indicates scaled expression across lineages. **f,** Lineage proportions (y-axis) versus serum troponin (x-axis) for pre-corticosteroid irMyocarditis samples. Unadjusted linear model p-values are shown. **Related to** Figure 1.

**Supplementary Figure 2.**
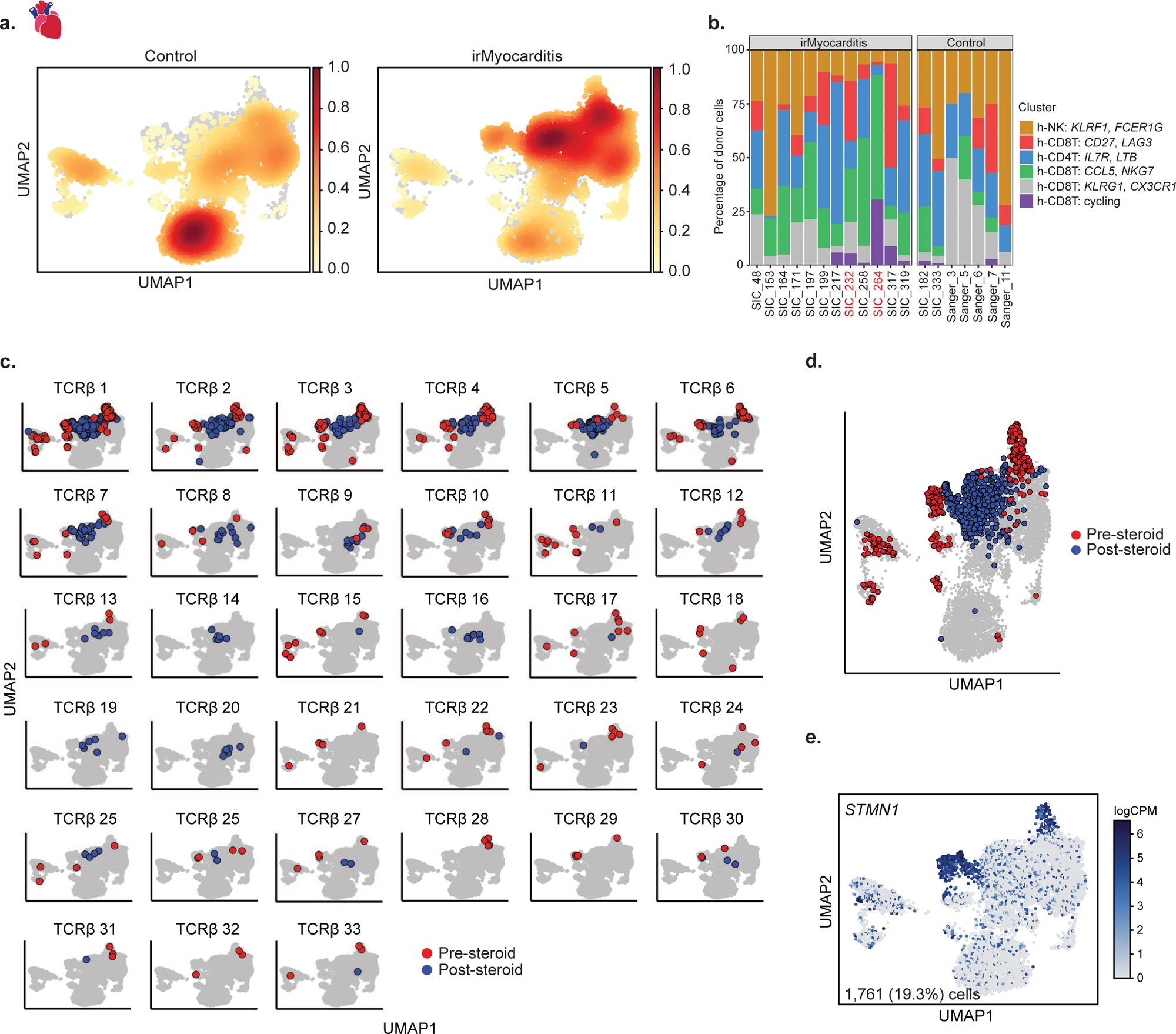
Lymphoid cells in irMyocarditis tissue. **a,** UMAP embedding of a cell density plot showing the contribution of cells from irMyocarditis cases and controls projected on the UMAP of T and NK cells derived from heart scRNA-seq data. **b,** Stacked bar plots showing the per-subset cellular composition per donor of each pre-corticosteroid or unenriched control sample. Red patient labels denote samples from patients with fatal irMyocarditis. **c, d** The 33 expanded TCR-β sequences (> 0.5% of TCR-β repertoire) from patient SIC_264 in T/NK UMAP space, color coded by whether the cell was found prior to (“pre-corticosteroid”, red) or after (“post-corticosteroid”, blue) administration of corticosteroids and second-line immunosuppression. Data shown by TCR-β clone (**c**) and in aggregate of all clones (**d**). **e,** Feature plot using color to indicate gene expression (logCPM) levels of *STMN1* projected onto the T/NK UMAP embedding. Cell numbers and percentages represent gene expression across all T/NK cells. **Related to** Figure 2.

**Supplementary Figure 3.**
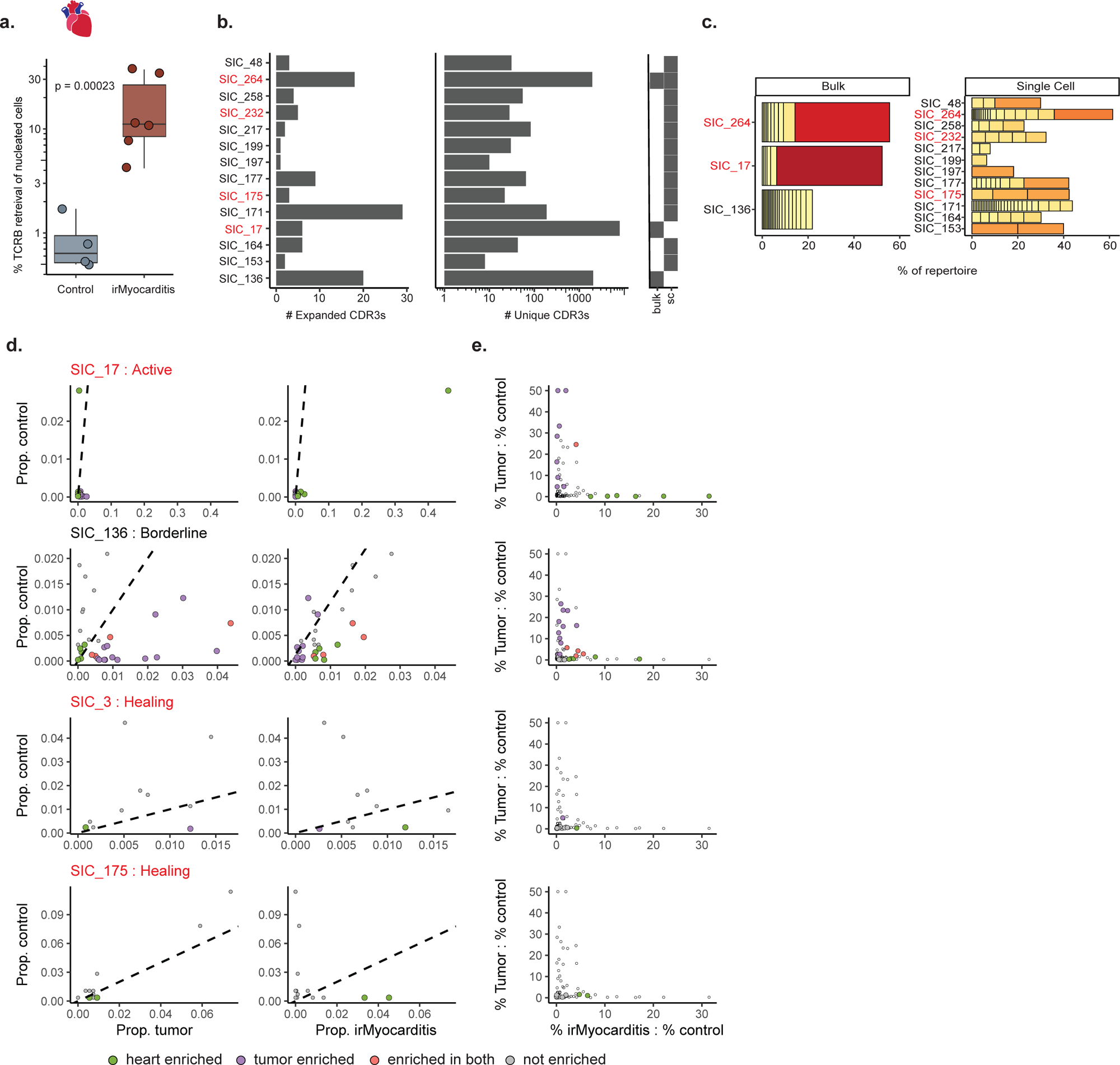
Expanded T-cell receptor β (TCR-β) sequences in irMyocarditis tissue. **a,** A boxplot showing the relative proportion of cells that recovered a TCR-β CDR3 sequence from marked areas of irMyocarditis (red) and control (blue) tissue; p = 0.0002, T-test (via Adaptive Biotechnologies). **b,** Expanded TCR-β CDR3 (left) and total TCR-β CDR3 sequences (right) recovered on a per patient basis from both scRNA-seq and bulk TCR-β CDR3 sequencing. **c,** Expanded TCR-β CDR3 from bulk sequencing (left) and scRNA-seq data from matched blood (right) in patients with irMyocarditis. **d,** Within each tissue type in each patient (“control”, “tumor”, or “irMyocarditis”), the frequency of each TCR-β clone is plotted on a per-patient basis and labeled by the pathological designation of the macroscopically dissected regions (SIC_17: Active; SIC_136: Borderline; SIC_3: Healing; SIC_175: Healing). Each point represents a TCR-β clone. The y-axis represents the proportion of a given TCR-β clone in the patient’s control tissue repertoire, and the x-axis represents the proportion of the TCR-β clone in their tumor TCR-β repertoire (left column) or irMyocarditis TCR-β repertoire (right column). Points are pseudocolored to represent a TCR-β clone that was expanded (> 0.5% of the heart or tumor repertoire) and enriched (Fisher’s exact test FDR < 5% compared to control) in heart (green), tumor (purple), both tissues (red), or neither tissue (grey). **e**, The frequency of each expanded TCR-β clone in heart and tumor tissue was calculated and then normalized by dividing by the frequency of the same clone in control tissue. Normalized TCR-β clone frequencies for heart (x-axis) and tumor tissue (y-axis) are plotted. Each plot shows the enriched TCR-β clones within each donor projected onto the aggregate data across all donors (Figure 3c-d). Individual data points from a given patient, representing TCR-β clones from that patient contributing to the aggregate plot, are colored by location of enrichment – heart (green), tumor (purple), both tissues (red), or not enriched (grey). In **a,** boxes represent the median (line) and interquartile range (IQR) with whiskers extending to the remainder of the distribution, no more than 1.5x IQR, with dots representing individual samples. Throughout the figure, red patient labels denote cases of fatal irMyocarditis. **Related to** Figure 3.

**Supplementary Figure 4.**
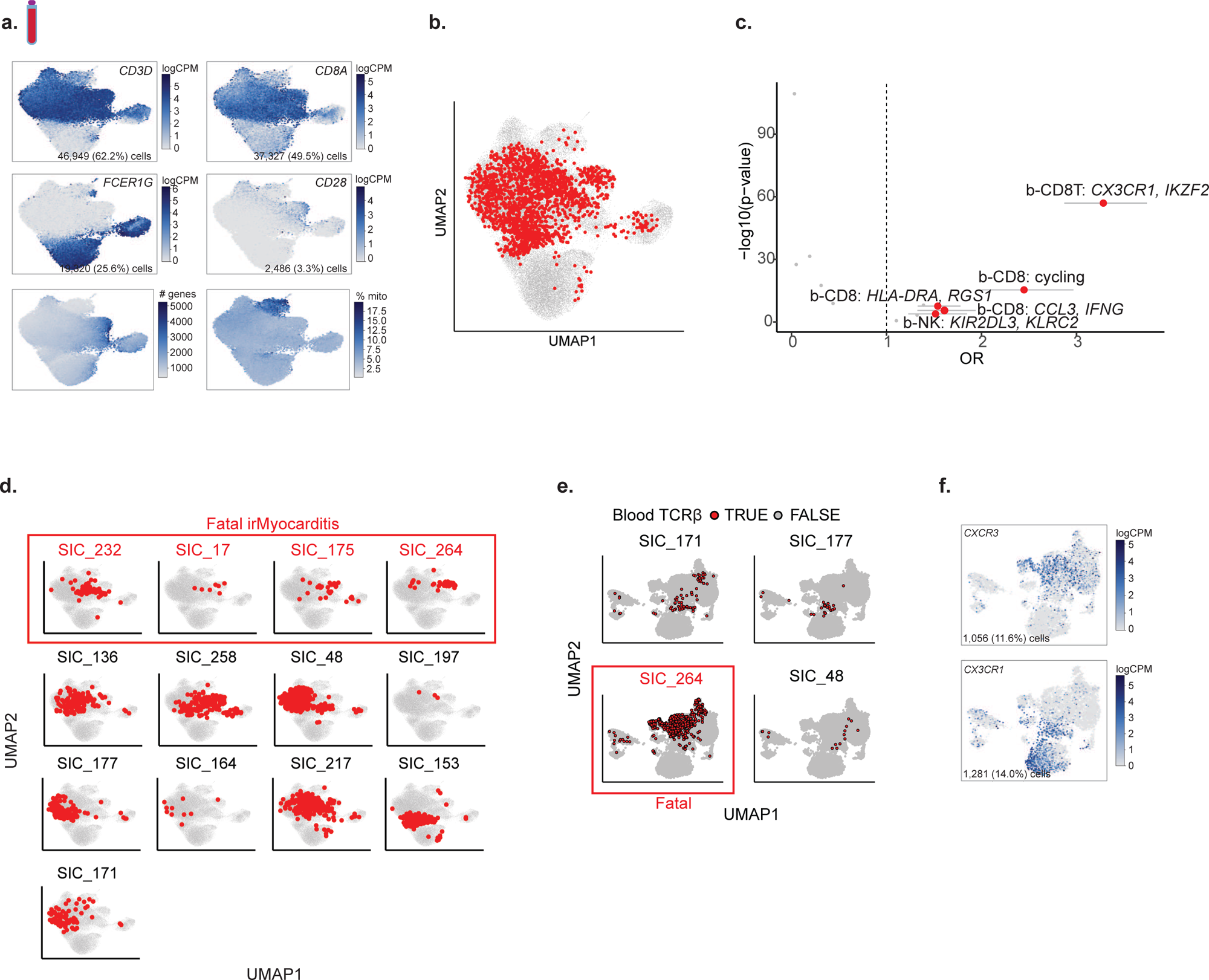
CD8 T and NK cells in blood. **a,** Feature plots using color to indicate gene expression (logCPM) levels of the indicated marker genes, number of unique genes expressed by each cells (bottom left) and percent mitochondrial genes (bottom right), projected onto the blood CD8 T/NK UMAP embedding. Cell numbers and percentages represent gene expression across all blood CD8 T/NK cells. **b,** Blood CD8 T/NK UMAP highlighting circulating cells (in red) that express a TCR-β sequence found to be expanded in irMyocarditis hearts (combined scRNA-seq and bulk TCR-β sequencing data). **c,** A volcano plot showing the results of a logistic regression model investigating the likelihood of a cell in a given CD8 T/NK cell subset in blood containing a TCR-β CDR3 sequence that was expanded in heart tissue. Red points denote cell subsets with statistically significant sharing (FDR < 0.05, likelihood-ratio test) **d**, Cells in blood for which the same TCR-β was expanded in paired irMyocarditis heart and blood sample from the same patient are shown in red on CD8 T/NK blood UMAP embeddings. Each patient with paired heart and blood samples is shown. **e,** UMAP of heart T and NK cells highlighting cells that express expanded TCR-β CDR3 sequences that were found in blood on a per-patient basis. **f,** Feature plots using color to indicate gene expression (logCPM) levels of the indicated genes projected onto the heart T and NK UMAP embedding. Cell numbers and percentages represent gene expression across all heart T and NK cells. In **e**, error bars represent 95% confidence intervals. In **f** and **g**, red patient labels denote cases of fatal irMyocarditis. **Related to** Figure 3.

**Supplementary Figure 5.**
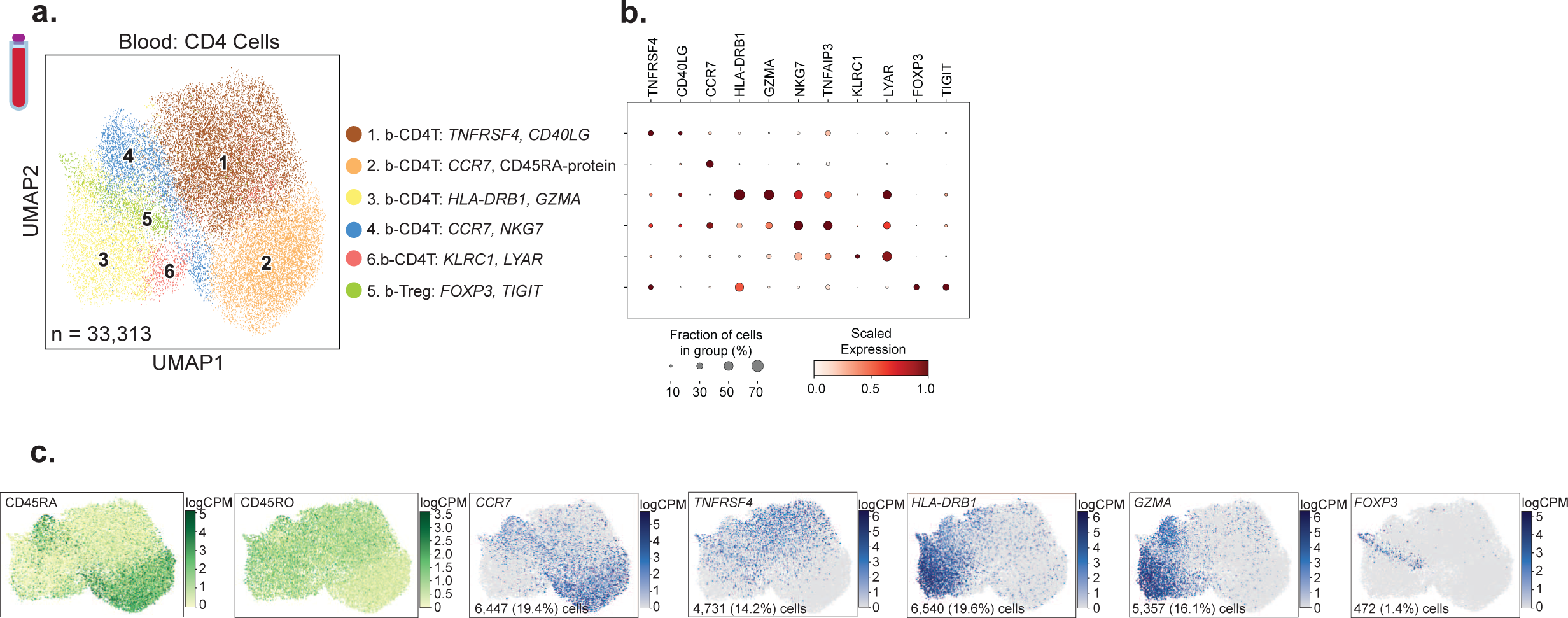
CD4 T cells in blood. **a,** UMAP embedding of 33,313 CD4 T cells from blood, colored by the six defined cell subsets labeled on the right. Cell subset number was assigned according to the absolute number of cells detected per subset. **b,** Dot plot showing top marker genes for each CD4 T cell subset in the blood. Dot size represents the percent of cells in the subset with non-zero expression of a given gene. Color indicates scaled expression across subsets. **c,** Left: feature plots using color to indicate surface protein levels (logCPM) of CD45RA and CD45RO protein (as determined by CITE-seq) projected onto the blood CD4 T cell UMAP embedding. Right: feature plots using color to indicate gene expression levels (logCPM) of the indicated genes projected onto the blood CD4 T cell UMAP embedding. Cell numbers and percentages represent gene expression across all blood CD4 T cells. **Related to** Figure 3.

**Supplementary Figure 6.**
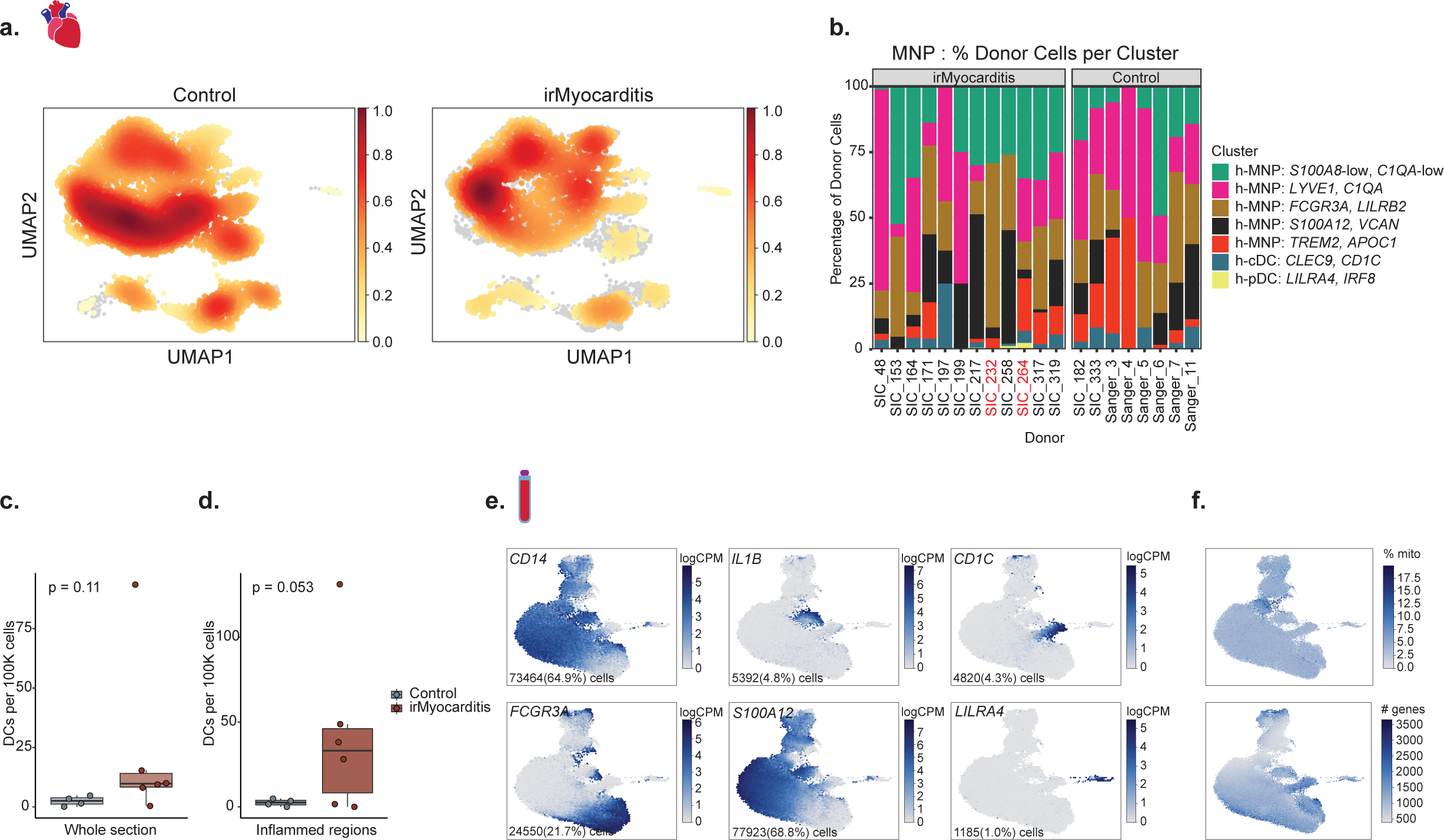
MNP populations in heart. **a,** Embedding of cell density plot showing the contribution of cells from irMyocarditis versus control samples projected on the UMAP of MNP cells derived from heart scRNA-seq data. **b**, Stacked bar chart depicting the relative contributions of cells in each MNP subset (colored coded on the right) on a per donor basis from each pre-corticosteroid irMyocarditis or unenriched control sample. **c**, Comparison of CD1c+ cell density measured by immunohistochemical staining of control heart sections (left column) versus whole slides from irMyocarditis heart sections (right column); p = 0.11, one-sided T-test. **d**, Comparison of CD1c+ cell density measured by immunohistochemical staining of control heart sections (left column) versus regions of inflammation in irMyocarditis heart sections (right column); p = 0.053, one-sided T-test. **e**, Feature plots using color to indicate gene expression (logCPM) levels of the indicated genes projected onto the blood MNP UMAP embedding. Cell numbers and percentages represent gene expression across all blood MNP cells. **f**, Feature plots showing percent mitochondrial genes (top panel), or number of unique genes expressed by each cell (bottom panel) on the UMAP of MNP cells derived from blood scRNA-seq data. **Related to** Figure 4.

**Supplementary Figure 7.**
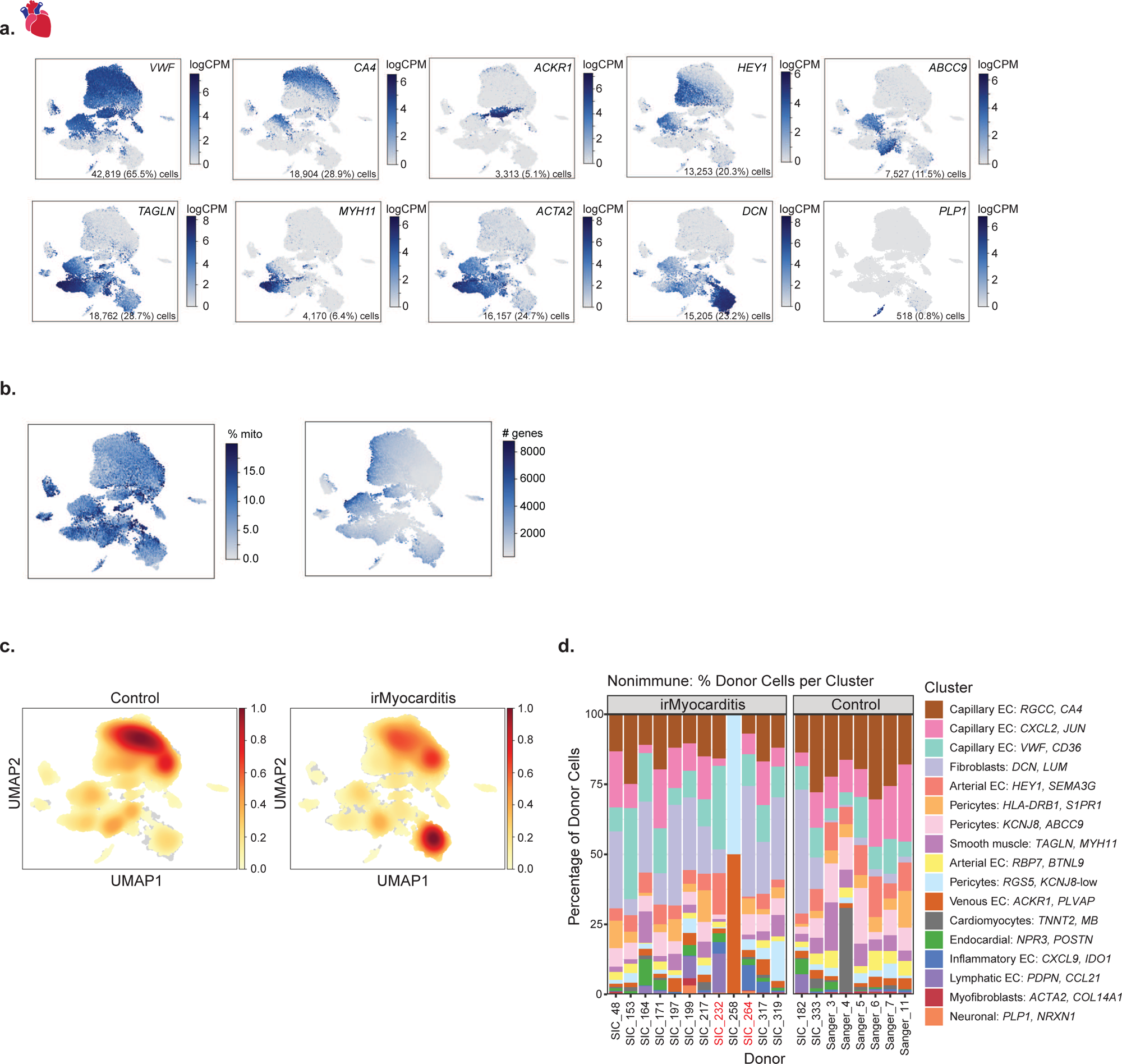
Non-immune populations in irMyocarditis heart tissue. **a,** Feature plots using color to indicate marker gene expression (logCPM) levels of the indicated genes projected onto the heart non-immune UMAP embedding. Cell numbers and percentages represent gene expression across all heart non-immune cells. **b**, Feature plots showing percent mitochondrial genes (left panel), or number of genes expressed by each cell (right panel) on the UMAP of non-immune cells derived from heart scRNA-seq data. **c**, UMAP embedding of cell density plot showing the contribution of cells from irMyocarditis versus control samples projected on the UMAP of non-immune cells derived from heart scRNA-seq data. **d**, Stacked bar chart depicting the relative contributions of cells in each non-immune cell subset (color coded to the right) from each pre-corticosteroid irMyocarditis or unenriched control sample. **Related to** Figure 5.

**Supplementary Figure 8.**
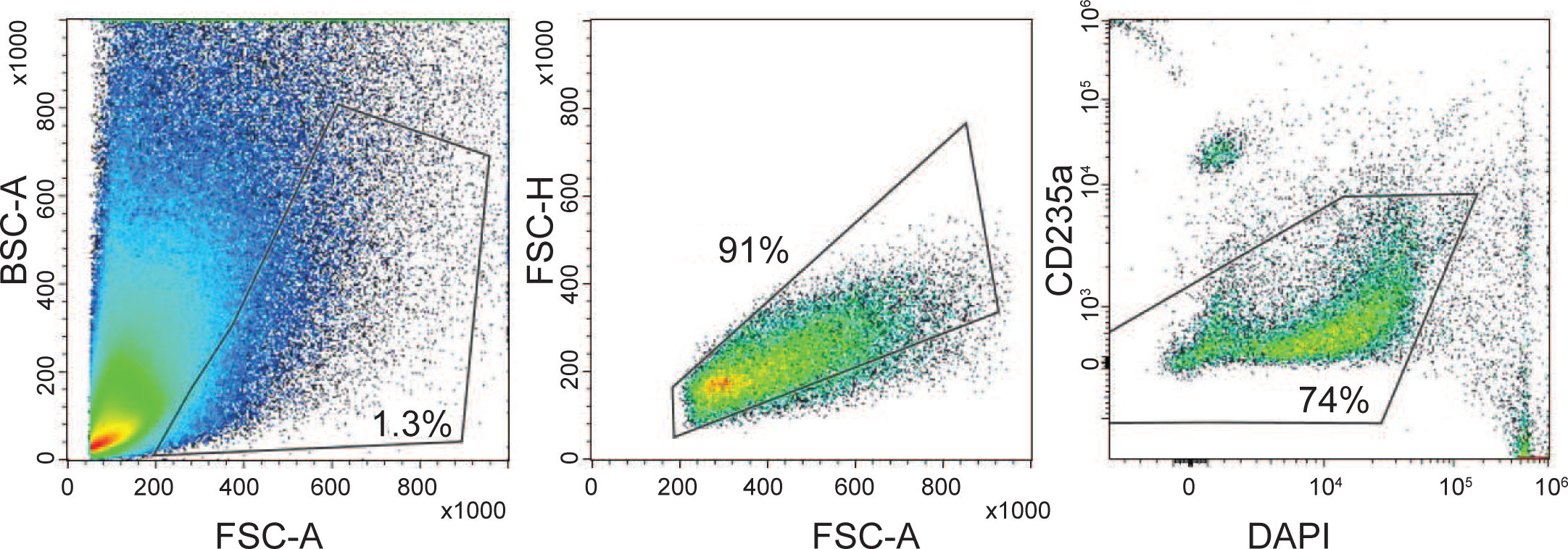
Sorting strategy for myocardial samples. Pseudocolor plots showing the applied sequential gating strategy to sort live cells for downstream scRNAseq from a representative myocardial sample. Numbers indicate the percentage within the indicated gate. DAPI^-^CD235a^-^ cells were collected for downstream analysis. **Related to Methods.**

